# Estimating the impact of differential adherence on the comparative effectiveness of stool-based colorectal cancer screening using the CRC-AIM microsimulation model

**DOI:** 10.1101/2020.08.30.271858

**Authors:** Andrew Piscitello, Leila Saoud, A. Mark Fendrick, Bijan J. Borah, Kristen Hassmiller Lich, Michael Matney, A. Burak Ozbay, Marcus Parton, Paul J. Limburg

## Abstract

**Background:** Real-world adherence to colorectal cancer (CRC) screening strategies is imperfect. The CRC-AIM microsimulation model was used to estimate the impact of imperfect adherence on the relative benefits and burdens of guideline-endorsed, stool-based screening strategies.

**Methods:** Predicted outcomes of multi-target stool DNA (mt-sDNA), fecal immunochemical tests (FIT), and high-sensitivity guaiac-based fecal occult blood tests (HSgFOBT) were simulated for 40-year-olds free of diagnosed CRC. For robustness, imperfect adherence was incorporated in multiple ways and with extensive sensitivity analysis. Analysis 1 assumed adherence from 0%-100%, in 10% increments. Analysis 2 longitudinally applied real-world first-round differential adherence rates (base-case imperfect rates=40% annual FIT vs 34% annual HSgFOBT vs 70% triennial mt-sDNA). Analysis 3 randomly assigned individuals to receive 1, 5, or 9 lifetime (9=100% adherence) mt-sDNA tests and 1, 5, or 9 to 26 (26=100% adherence) FIT tests. Outcomes are reported per 1000 individuals compared with no screening.

**Results:** Each screening strategy decreased CRC incidence and mortality versus no screening. In individuals screened between ages 50-75 and adherence ranging from 10%-100%, the life-years gained (LYG) for triennial mt-sDNA ranged from 133.1-300.0, for annual FIT from 96.3-318.1, and for annual HSgFOBT from 99.8-320.6. At base-case imperfect adherence rates, mt-sDNA resulted in 19.1% more LYG versus FIT, 25.4% more LYG versus HSgFOBT, and generally had preferable efficiency ratios while offering the most LYG. Completion of at least 21 FIT tests is needed to reach approximately the same LYG achieved with 9 mt-sDNA tests.

**Conclusions:** Adherence assumptions affect the conclusions of CRC screening microsimulations that are used to inform CRC screening guidelines. LYG from FIT and HSgFOBT are more sensitive to changes in adherence assumptions than mt-sDNA because they require more tests be completed for equivalent benefit. At imperfect adherence rates, mt-sDNA provides more LYG than FIT or HSgFOBT at an acceptable tradeoff in screening burden.

## Introduction

Screening and subsequent treatment of pre-symptomatic neoplasia has been shown to reduce the incidence and mortality of colorectal cancer (CRC).[1, 2] Widely endorsed CRC screening modalities include structural examinations (e.g., colonoscopy, computed tomography colonography [CTC], and flexible sigmoidoscopy [SIG]) and stool-based tests (e.g., fecal immunochemical testing [FIT], high-sensitivity guaiac-based fecal occult blood tests [HSgFOBT], and multitarget stool DNA [mt-sDNA]). Ideally, randomized clinical trials (RCTs) would be conducted to evaluate the comparative effectiveness of different CRC screening strategies. However, obtaining long-term clinical trial data for all the various permutations of CRC screening strategies (i.e. screening modality, start age, stop age, and interval) is not feasible. Therefore, outcomes of microsimulation models have been used by clinical societies and expert panels (i.e., American Cancer Society [ACS] and US Preventive Services Task Force [USPSTF]), in conjunction with systematic evidence reviews, to determine appropriate CRC screening strategies.[3, 4] Three CRC microsimulation models have been independently developed with funding from the Cancer Intervention and Surveillance Modeling Network (CISNET) Colorectal Working Group.[5–8] In 2016, CISNET published a set of model-recommendable screening strategies that were based on a combination of modeling results and consultation with USPSTF members.[4]

The CISNET analyses used to inform USPSTF (and other) guidelines assume perfect (100%) adherence with all CRC screening, follow-up, and surveillance procedures over each individual’s lifetime.[4, 9] Therefore, the recommendations are based on what is theoretically, rather than practically, achievable. Although assuming perfect adherence to all screening strategies provides the opportunity for impartial comparison across the analyzed modalities, this assumption inherently limits real-world application of the estimated outcomes. Indeed, perfect adherence has been recognized by the CISNET group and others as a limitation to traditional simulation analyses.[9–11] Although currently sparse, real-world data referent to longitudinal CRC screening demonstrate variable participation rates both across and within populations, with none approaching the modeled assumption of 100%.[12–15] To more realistically assess the relative benefits and burdens of stool-based screening in microsimulation modeling, imperfect adherence needs to be taken into consideration.

A new CRC microsimulation model, Colorectal Cancer and Adenoma Incidence and Mortality Microsimulation Model (CRC-AIM), was developed based on previously reported parameters of CRC-SPIN.[16, 17] Because of the substantial differences among screening modalities (e.g. bowel prep, invasiveness, time off work, adverse events), this analysis is limited to stool tests, which have similar burdens and no direct complications. The objective of the current analyses was to use CRC-AIM to estimate the impact of imperfect adherence on the relative benefits and burdens of guideline-endorsed stool-based screening strategies.

## Methods

### Microsimulation model

CRC-AIM natural history, which is based on a published methodology,[18–20] is used to model the sequence of adenoma to carcinoma progression in unscreened patients, as briefly described herein. As individuals age, the risk of adenomas generally increases. Adenomas may grow and transition to preclinical cancer, which may transition to symptomatic CRC. CRC screening facilitates the removal of adenomas and potential early detection of preclinical CRC.[4] The ability of a stool-based CRC screening test to detect an adenoma or preclinical CRC is dependent on test performance (i.e., sensitivity) and test completion (i.e., adherence). Positive screening tests are followed-up with a diagnostic (i.e. follow-up) colonoscopy. Further description of the natural history and screening components of the CRC-AIM model is provided in the **Supplemental Material**. Full details of the natural history and screening components, as well as validation results evaluating qualitative and quantitative outputs to those from CRC-SPIN, SimCRC, and MISCAN,[4, 9] are available elsewhere.[17]

For the primary analyses, all CRC screening test performance assumptions (sensitivity, specificity, and complications) were identical to the CISNET modeling analyses used to inform the 2016 USPSTF guideline recommendations (**Table 1 and Supplemental Fig S1**).[4, 9]

**Table 1.** Screening test characteristic inputs. Reproduced and adapted with permission from Knudsen et al, 2016.[4]

|  | Screening Test, Source Citation for Input |  |  |  |
| --- | --- | --- | --- | --- |
| Screening test characteristic | Colonoscopy (within reach, per lesion)<sup>a</sup> | FIT (per person) | HSgFOBT (per person) | mt-sDNA (per person) |
| Sensitivity for CRC, % | 95<sup>b</sup> | 73.8[21] | 70[22] | 92.3[21] |
| Sensitivity for adenomas $\geq 10$ mm, % | 95[23] | 23.8<sup>c</sup>, [21] | 23.9[22] | 42.4<sup>c</sup>, [21] |
| Sensitivity for adenomas 6-9 mm, % | 85[23] | 7.6<sup>d</sup>, [21] | 12.4[22] | 17.2<sup>d</sup>, [21] |
| Sensitivity for adenomas 1-5 mm, % | 75[23] |  | 7.5<sup>e</sup>, [22] |  |
| Specificity, % | 86<sup>f</sup>, [24] | 96.4[21] | 92.5[22] | 89.8[21] |
| Reach, % | 95 to end of cecum, remainder between rectum and cecum<sup>b,g</sup> | Whole colorectum<sup>b</sup> | Whole colorectum[22] | Whole colorectum<sup>b</sup> |
| Risk of complications (serious GI, other GI, and CV complications) | Age-specific risks<sup>h</sup>, [25-27] | 0[28] | 0[28] | 0[28] |

COL, colonoscopy; CRC, colorectal cancer; CV, cardiovascular; FIT, fecal immunochemical test with a positivity cutoff of $\geq 100$ ng of hemoglobin (Hb) per mL of buffer ( $\geq 20$ mcg Hb/g of feces); GI, gastrointestinal; HSgFOBT, high-sensitivity guaiac-based fecal occult blood test; mt-sDNA, multitarget stool DNA test.
<sup>a</sup>It was assumed that the same test characteristics for screening colonoscopies applied to colonoscopies for diagnostic follow-up or for surveillance.
<sup>b</sup>No source, input is by assumption.
<sup>c</sup>Sensitivity for persons with advanced adenomas (ie, adenomas $\geq 10$ mm or adenomas with advanced histology). Sensitivity was not reported for the subset of persons with $\geq 10$ mm adenomas.
<sup>d</sup>Sensitivity for persons with nonadvanced adenomas.
<sup>e</sup>It was assumed that 1–5 mm adenomas do not bleed and therefore cannot cause a positive stool test. It was also assumed that HSgFOBT can be positive because of bleeding from other causes, the probability of which is equal to positivity rate in persons without adenomas.
<sup>f</sup>The lack of specificity with endoscopy reflects the detection of nonadenomatous polyps, which, in the case of colonoscopy, leads to unnecessary polypectomy, which is associated with an increased risk of colonoscopy complications.
<sup>g</sup>Full reach (to the cecum) is assumed to be achieved 95% of the time and if reach is only partial, a second colonoscopy is performed.[4]
<sup>h</sup>See **Fig S1** in the Supplement for details on age-specific risks.

### CRC screening outcomes

Predicted CRC screening outcomes were simulated for 4 million individuals born in 1975. Efficacy was measured by number of life-years gained (LYG) compared with no CRC screening. Number of colonoscopies was used as a proxy for burden and harms. The number of total stool tests, complications from colonoscopies, CRC cases, CRC deaths, life-years with CRC, incidence reduction, and mortality reduction were analyzed as additional outcomes. All outcomes were reported per 1,000 individuals free of diagnosed CRC at age 40.

For each stool-based screening modality, up to 18 strategies were evaluated, consisting of a unique combination of screen interval (1, 2 y FIT/HSgFOBT, 2, 3 y mt-sDNA), age to begin screening (45, 50, 55 y), and age to end screening (75, 80, 85 y) (**Table 2**). Detailed screening outcomes are presented from selected stool-based screening strategies, namely annual FIT, annual HSgFOBT, and triennial mt-sDNA for individuals screened between ages 50–75 and 45–75, since these are the strong and qualified USPSTF and American Cancer Society intervals and age recommendations.[3, 4] Screening outcomes from other stop-start age combinations can be accessed at https://github.com/CRCAIM/CRC-AIM-Public.

**Table 2.**
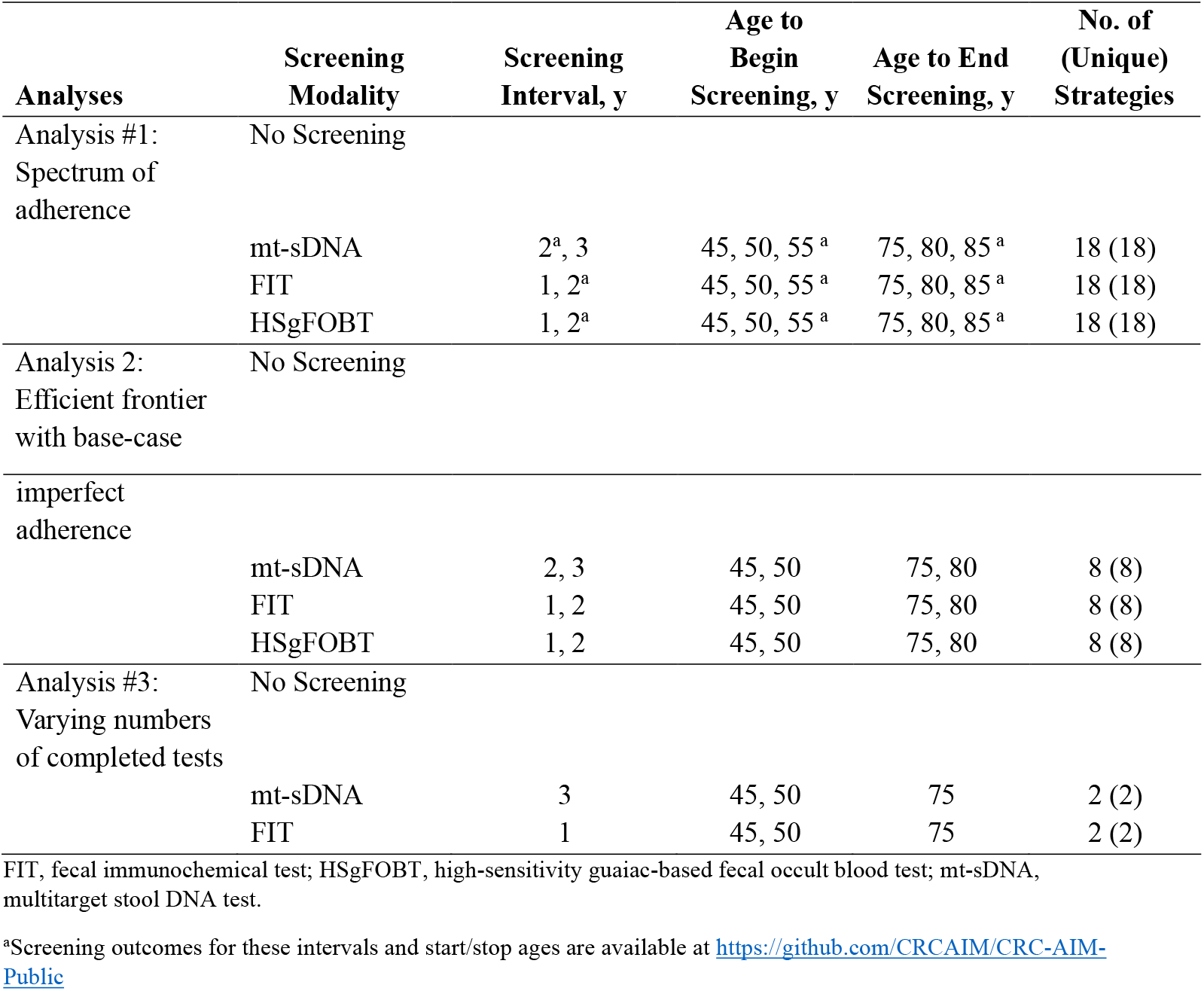
Screening strategies evaluated for each adherence analysis.

### Adherence analyses

Modeling of real-world CRC screening adherence is complicated and, at present, there is no universally accepted methodology. To sufficiently evaluate the impact of adherence on CRC screening outcomes, multiple adherence analyses were performed, described below. Since the purpose of these analyses was to determine the impact of adherence only on stool-based screening, perfect adherence to diagnostic and surveillance colonoscopies was assumed for all analyses in accordance with CISNET models.[4, 9]

#### Analysis 1

The purpose of analysis 1 was to evaluate the comparative effectiveness of stool-based testing screening strategies assuming a spectrum of adherence rates for each modality. Adherence was set by assuming a fixed annual likelihood to comply with each stool-based screening strategy ranging from 0% to 100%, in 10% increments. It was assumed that individuals were offered a stool-based test every year unless they were not due for screening.

#### Analysis 2

The purpose of Analysis 2 was to estimate the comparative impact of differential adherence to each stool test using real-world first-round screening participation rates and applying them as a fixed annual likelihood to comply. Due to the heterogeneity of reported real-world adherence, several different imperfect adherence scenarios were evaluated. The base-case imperfect adherence rates were 40% FIT[29] vs 34% HSgFOBT[29] vs 70% mt-sDNA[30]. Adherence of 40% for FIT was loosely estimated based on pooling participation rates reported in a meta-analysis[29] resulting in 42% adherence (range of individual studies was 26% to 62%). Adherence for HSgFOBT was found to be approximately 16% lower relative to FIT in a meta-analysis,[29] and therefore was analyzed as 34% (40% FIT / 1.16 =34%). Adherence of 70% for mt-sDNA was based on a retrospective cohort analysis.[30]

##### Efficient frontier of stool-based screening strategies

Using a similar methodology to the CISNET Colorectal Cancer Working Group (CWG),[4] efficient frontiers were used to visualize tradeoffs between benefits and burdens, and tables were used to summarize results. Two different sets of benefit/burden pairs were evaluated. The first set of outcomes for efficient frontier generation used LYG and the number of required colonoscopies. The second set of outcomes used LYG and patient hours related to the screening process. Based on previously published CISNET assumptions, patient time related to the screening process was assumed to be 16 hours for colonoscopy, 112 hours for complications due to colonoscopy, and 1 hour for FIT.[31] Patient time to complete mt-sDNA was assumed to be the same as FIT and for HSgFOBT was assumed to be 2 hours: ½ hour was assumed for each of the 3 evacuation/collection/storage processes required for HSgFOBT plus ½ hour for instructions/unboxing/setup/mailing.

Each benefit/burden pair were plotted to generate efficient frontiers. Strongly dominated strategies (i.e, strategies that required more colonoscopies or patient hours for fewer LYG) were discarded. The incremental number of LYG per 1000 (ΔLYG) and incremental number of colonoscopies per 1000 (ΔCOL) or patient hours related to the screening process (Δpatient hours) were computed. The efficiency ratio (ΔCOL/ΔLYG or Δpatient hours/ΔLYG) for each remaining strategy was calculated. Strategies with fewer LYG but a higher efficiency ratio than another strategy were discarded as weakly dominated. The efficient frontier was the line that connected the efficient strategies; strategies that had LYG within 98% of the efficient frontier were considered near-efficient. Typically, a recommendable strategy must be efficient or near efficient, offer LYG within 90% of a benchmark COL strategy, must have an efficiency ratio (e.g. incremental COL per incremental LYG) at or below the range of CISNET model-derived acceptable thresholds, based on the selected benchmark COL strategy (e.g. incremental ratios between 39-65 for a 10-year benchmark COL when the screening window is 50–75), and, if all criteria are met, offer the most LYG.[4] However, in the context of this analysis, %LYG of the benchmark COL is not applicable since it is assumed that real-world adherence is imperfect (including that of the benchmark COL, which is not evaluated), and for simplicity, we assume efficiency ratio thresholds of 39-65 irrespective of a benchmark COL strategy. Efficient frontier plots for all strategies were generated for perfect adherence rates (to replicate CISNET results[4, 9]) and imperfect adherence rates. Calculated frontier outcomes and efficiency ratios were then tabulated for all strategies, from which a subset could be taken to assess outcomes for a particular screen window.[4, 9] Mimicking an alternate approach to this algorithm,[32] frontier outcomes were separately calculated for all modalities for a given screen window (e.g., 50-75, 45-75), as opposed to including all screen windows on the same frontier prior to calculating frontier outcomes,[4, 9] since it is presumed a priori that only one start-stop age pair would be implemented in practice.

#### Analysis 3

The purpose of analysis 3 was to evaluate the comparative effectiveness of stool-based testing assuming varying numbers of completed stool-based tests. This analysis represents individuals who are randomly adherent during a screening window from ages 50-75. Simulated individuals were randomly assigned to up to 1, 5, or 9 (9=100% adherence) mt-sDNA tests and up to 1, 5, or 9 to 26 (26=100% adherence) FIT tests during the lifetime screening window. Simulated individuals may have completed fewer than their assigned tests if they died or had a positive stool test. Only recommended screening strategies of triennial mt-sDNA and annual FIT were included in this analysis; therefore, no efficient frontiers were generated.[3] An additional analysis assuming a screening window from ages 45-75 was also performed, with a corresponding set of randomly completed tests (e.g., 31=100% adherence for FIT and 11=100% adherence for mt-sDNA).

### Sensitivity analysis

#### Test performance sensitivity analyses

The same screening characteristics for FIT, mt-sDNA, and HSgFOBT were used in the CRC-AIM primary analyses and the CISNET models.[4, 9] These screening characteristics for FIT and mt-sDNA were derived from data generated in a cross-sectional study of 9989 participants (aged 50-84 years) at average risk of CRC who underwent colonoscopy, mt-sDNA, and FIT screening (DeeP-C study; clinicaltrials.gov identifier, NCT01397747).[21] Adenoma findings in the published report by Imperiale et al[21] were not distinguished by size or location, but rather were categorized as advanced (≥10 mm) or non-advanced adenomas. Thus, the CISNET models and the CRC-AIM model primary analyses used the published sensitivity of advanced adenomas as a proxy for the sensitivity of adenomas ≥10 mm and non-advanced adenomas as a proxy for the sensitivity of adenomas 1-5 mm and 6-9 mm combined.[9] Additional outcomes for adherence analyses #1 and #3 were generated using screening characteristics for FIT and mt-sDNA derived from more granular estimates of diagnostic sensitivity, stratified by adenoma size and location (rectal, distal, proximal), and age-based specificity data collected in DeeP-C[21] to which the study sponsor had access. The screening characteristics for the sensitivity analysis are shown in **Table 3**. All other aspects of the CRC-AIM modeling in the sensitivity analyses were the same as the primary analyses. Finally, for informational purposes, sensitivity to detect adenomas based on size alone for FIT and mt-sDNA, derived from DeeP-C (clinicaltrials.gov identifier, NCT01397747),[21] are in **Supplemental Table S1**.

**Table 3.** Screening characteristics for Deep-C (clinicaltrials.gov identifier, NCT01397747) sensitivity analyses using granular adenoma size and location data.

|  | Sensitivity |  |  |  |  |  |  |  |  |  | Specificity |  |  |  |  |
| --- | --- | --- | --- | --- | --- | --- | --- | --- | --- | --- | --- | --- | --- | --- | --- |
|  | Adenomas, mm |  |  |  |  |  |  |  |  |  |  |  |  |  |  |
|  | Rectal |  |  | Distal |  |  | Proximal |  |  | Cancer Stage | Age, y |  |  |  |  |
|  | ≤5 | 6-9 | ≥10 | ≤5 | 6-9 | ≥10 | ≤5 | 6-9 | ≥10 | I-IV | <60 | 60-64 | 65-69 | 70-74 | 75+ |
| FIT, % (n/N) | 3.3% (5/150) | 5.6% (5/89) | 30.1% (22/73) | 7.0% (39/558) | 21.5% (68/317) | 39.3% (83/211) | 5.9% (73/1236) | 6.6% (40/609) | 16.0% (65/406) | 73.8% (48/65) | 97.8% (1361/1392) | 96.5% (355/368) | 95.6% (1483/1551) | 96.7% (697/721) | 93.9% (399/425) |
| 95% CI | 1.1% - 7.6% | 1.8% - 12.6% | 19.9% - 42.0% | 5.0% - 9.4% | 17.1% - 26.4% | 32.7% - 46.3% | 4.7% - 7.4% | 4.7% - 8.8% | 12.6% - 19.9% | 61.5% - 84.0% | 96.9% - 98.5% | 94.0% - 98.1% | 94.5% - 96.6% | 95.1% - 97.9% | 91.2% - 96.0% |
| mt-sDNA, % (n/N) | 9.3% (14/150) | 21.3% (19/89) | 56.2% (41/73) | 14.5% (81/558) | 27.8% (88/317) | 57.3% (121/211) | 15.8% (195/1236) | 19.9% (121/609) | 34.0% (138/406) | 92.3% (60/65) | 94.4% (1314/1392) | 92.4% (340/368) | 89.1% (1382/1551) | 86.1% (621/721) | 81.2% (345/425) |
| 95% CI | 5.2% - 15.2% | 13.4% - 31.3% | 44.1% - 67.8% | 11.7% - 17.7% | 22.9% - 33.0% | 50.4% - 64.1% | 13.8% - 17.9% | 16.8% - 23.3% | 29.4% - 38.8% | 83.0% - 97.5% | 93.1% - 95.5% | 89.2% - 94.9% | 87.4% - 90.6% | 83.4% - 88.6% | 77.1% - 84.8% |

FIT, fecal immunochemical test; mt-sDNA, multitarget stool DNA test.
2-sided 95% confidence intervals calculated using the Exact (Clopper-Pearson) method.

#### Efficiency frontier sensitivity analyses

Sensitivity analyses of efficient frontier plots (analysis #2) were conducted using 50% FIT vs 43% HSgFOBT vs 70% mt-sDNA adherence and 60% FIT vs 52% HSgFOBT vs 70% mt-sDNA adherence based on existing data referent to test-specific adherence rates.[14, 33–38]

An additional sensitivity analysis was performed to evaluate an alternative approach to modeling imperfect adherence. CISNET has modeled imperfect adherence by splitting the population between those that are 100% adherent to screening versus those who are 0% adherent to screening (CISNET fixed split adherence approach).[39–41] This approach was replicated to evaluate its impact on base-case adherence assumptions for analysis #2.

## Results

### Analysis 1: Spectrum of adherence

Analysis #1 assumed a spectrum of adherence in 10% increments for stool-based screening strategies. Each screening strategy reduced CRC-related incidence and mortality compared with no screening. When the screening window was between ages 50-75, for triennial mt-sDNA, the LYG from 10% to 100% adherence ranged from 133.1 to 300.0, for annual FIT ranged from 96.3 to 318.1, and for annual HSgFOBT ranged from 99.8 to 320.6 (**Supplemental Table S2**). LYG for FIT was more sensitive to per-unit change in adherence rates ([318.1-96.3]/[100%-10%]=2.5 LYG/unit change) than mt-sDNA (1.9 LYG/unit change). LYG for HSgFOBT was also more sensitive to per-unit change (2.5 LYG/unit change) versus mt-sDNA. A matrix of relative LYG comparisons between mt-sDNA and FIT for all adherence combinations (**Fig 1**) illustrates regions of differential adherence where the tests have similar (<10%) or dissimilar (≥10%) differences in LYG.

**Fig 1.**
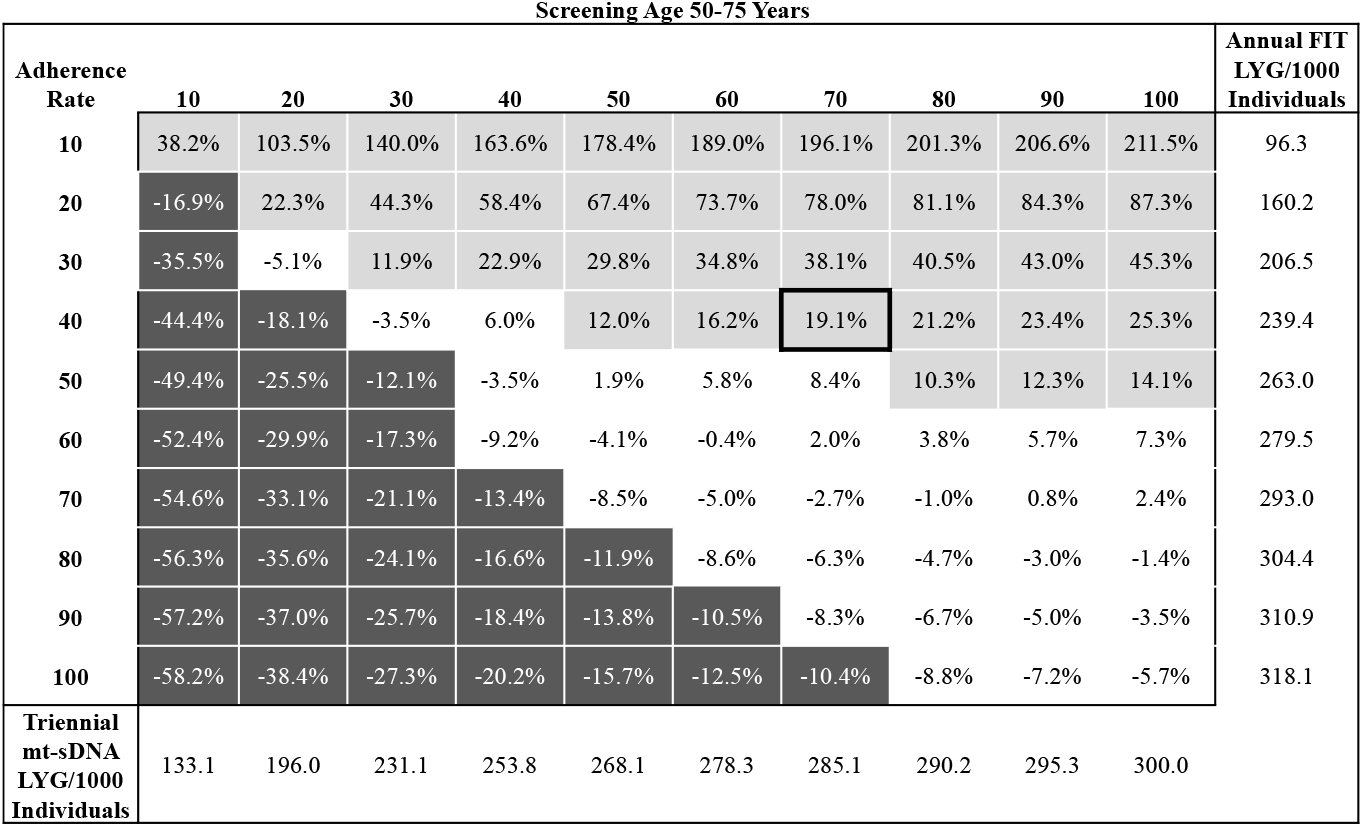
Percent difference in predicted life-years gained (LYG) per 1000 individuals by adherence rate for triennial multitarget stool DNA (mt-sDNA) versus annual fecal immunochemical test (FIT) in individuals free of diagnosed colorectal cancer at age 40 and screened between ages 50–75 years. White boxes indicate <10% difference between tests. Light gray boxes indicate ≥10% positive difference with mt-sDNA versus FIT. Dark gray boxes indicate ≥10% negative difference with mt-sDNA versus FIT. Outlined box indicates base-case imperfect adherence rates.

At perfect adherence, the LYG and reductions in CRC-related incidence and mortality were highest for annual HSgFOBT, followed by annual FIT and triennial mt-sDNA (**Fig 2**). The total number of stool tests was lower with mt-sDNA vs FIT and HSgFOBT (**Fig 2**). Total required colonoscopies was similar between mt-sDNA and FIT and was slightly higher for HSgFOBT (**Fig 2**). Other screening outcomes also vary across the spectrum of adherence assumptions and assumed screening windows (**Supplemental Table S2 and Supplemental Fig S2**).

**Fig 2.**
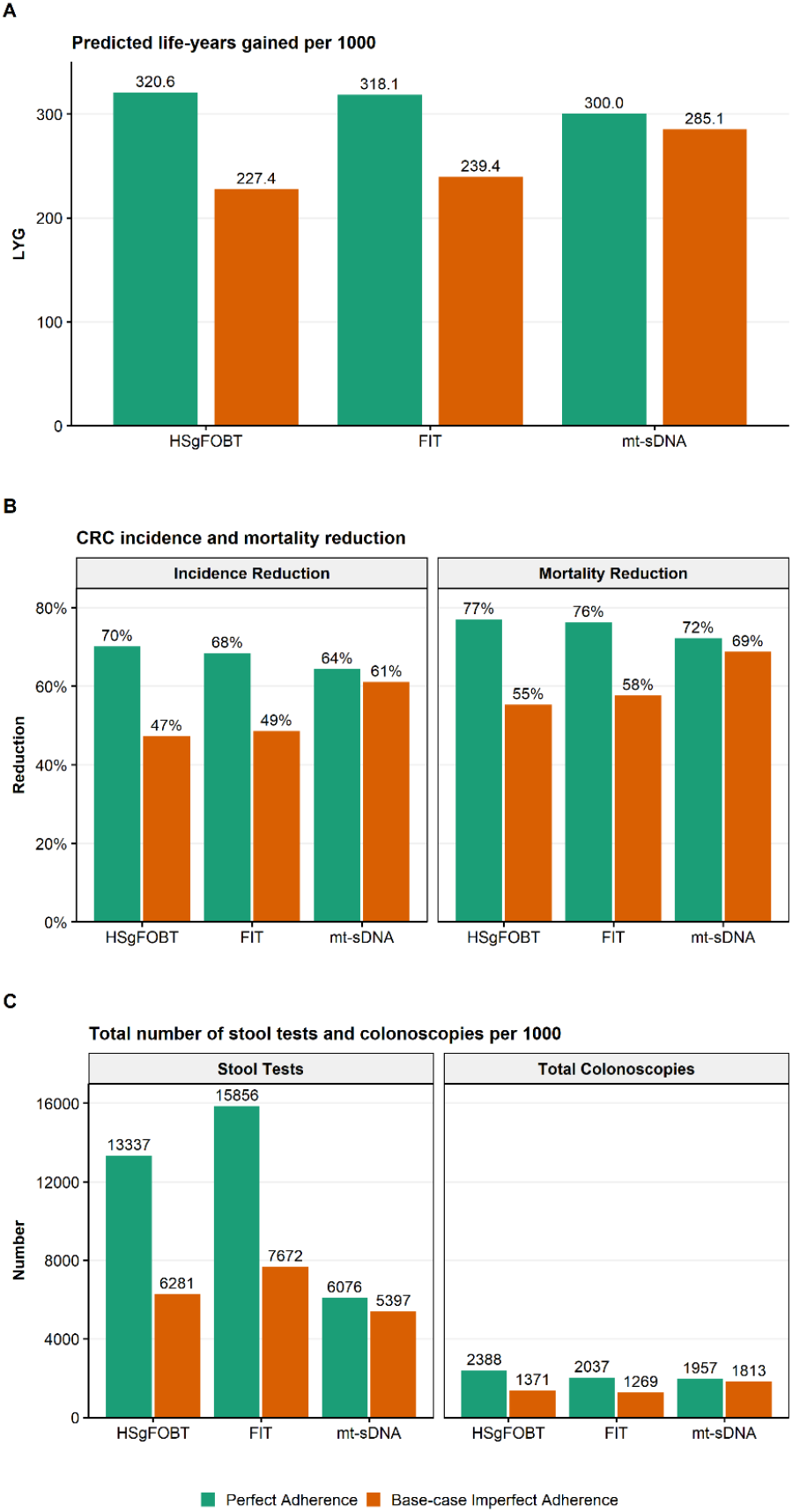
A) Predicted life-years gained (LYG), B) CRC-related incidence and mortality reduction, and C) total stool tests and colonoscopies (COL) per 1000 individuals screened from ages 50–75 compared with no screening assuming perfect (100%) adherence rates to annual FIT, annual HSgFOBT, and triennial mt-sDNA or base-case imperfect adherence (40% FIT vs 34% HSgFOBT vs 70% mt-sDNA).

### Analysis 2: Base-case imperfect adherence

At base-case imperfect adherence rates (40% annual FIT vs 34% annual HSgFOBT vs 70% triennial mt-sDNA) in individuals screened between ages 50–75, the number of LYG was highest for mt-sDNA resulting in 19.1% more LYG (285.1) versus FIT (239.4) and 25.4% more LYG versus HSgFOBT (227.4; **Fig 2**). Mt-sDNA had the highest number of colonoscopies, whereas FIT had the highest number of stool tests (**Fig 2**). A similar pattern of results was observed with a 45-75 screening window (**Supplemental Fig S3**).

#### Efficient frontier of stool-based screening strategies

The efficient frontiers by LYG relative to number of colonoscopies of stool-based screening strategies assuming perfect adherence or base-case imperfect adherence rates are shown in **Fig 3**. When shifting from perfect to base-case imperfect adherence rates, strategies that are on or off the efficient frontier can change, and efficiency ratios can substantially change. (**Supplemental Tables S3 and S4**). Adherence-related changes to efficient frontiers impact model-recommendable strategies. When the screen window is fixed at 50-75 followed by recalculating efficiency comparisons, the perfect adherence model-recommended strategy is annual FIT (incremental ratio 12.5 COL/LYG, as opposed to near-efficient biennial mt-sDNA and efficient annual HSgFOBT, which offer more LYG but exceed the efficiency ratio [slope] threshold of 39-65 at incremental ratios of 196.6 COL/LYG and 142.8 COL/LYG, respectively) and the base-case imperfect adherence recommended strategy is biennial mt-sDNA (incremental ratio of 12.8 COL/LYG, with no other strategies offering more LYG) (**Supplemental Table S5**). Similar results were obtained for a screening window from 45-75 (**Supplemental Table S5**).

**Fig 3.**
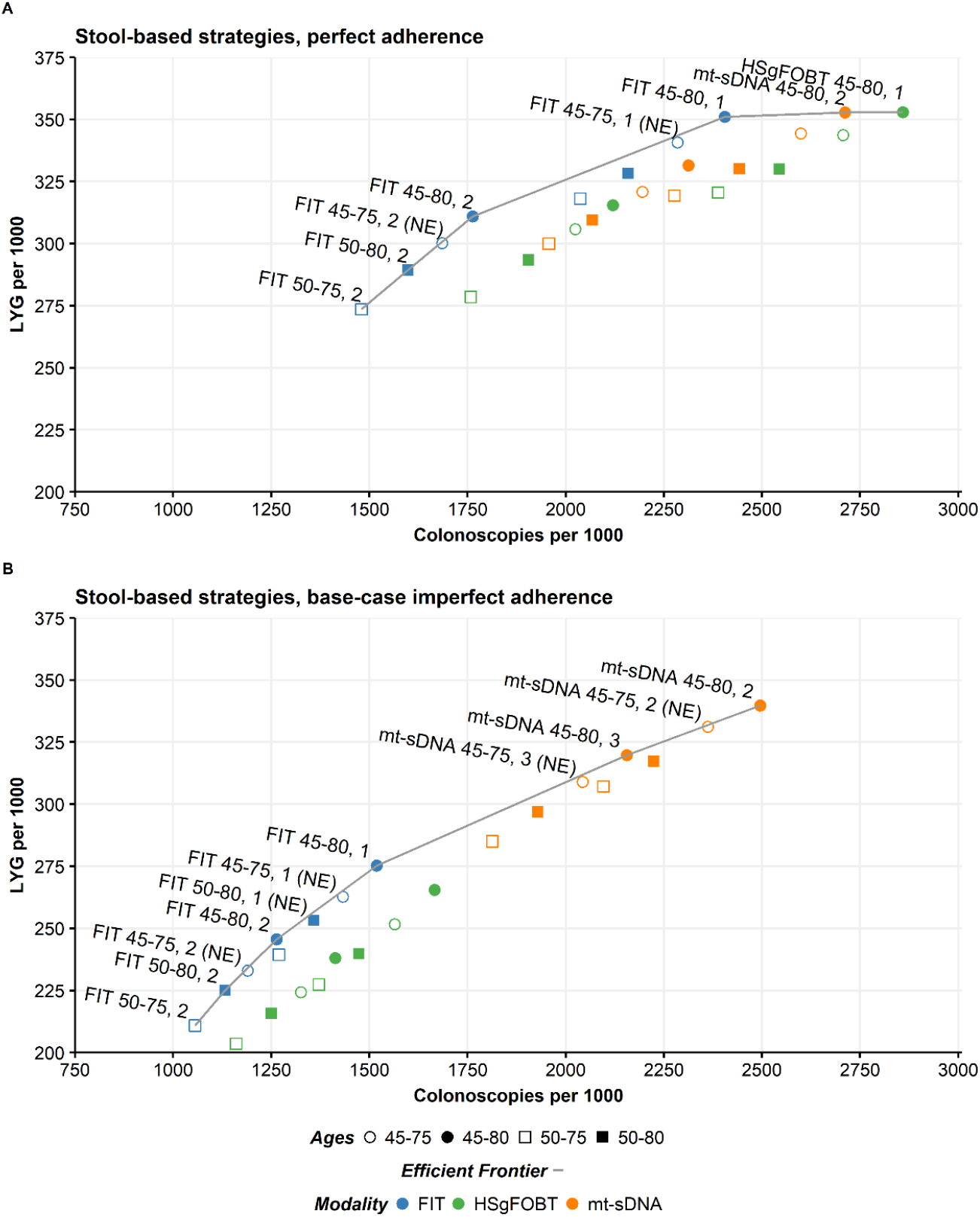
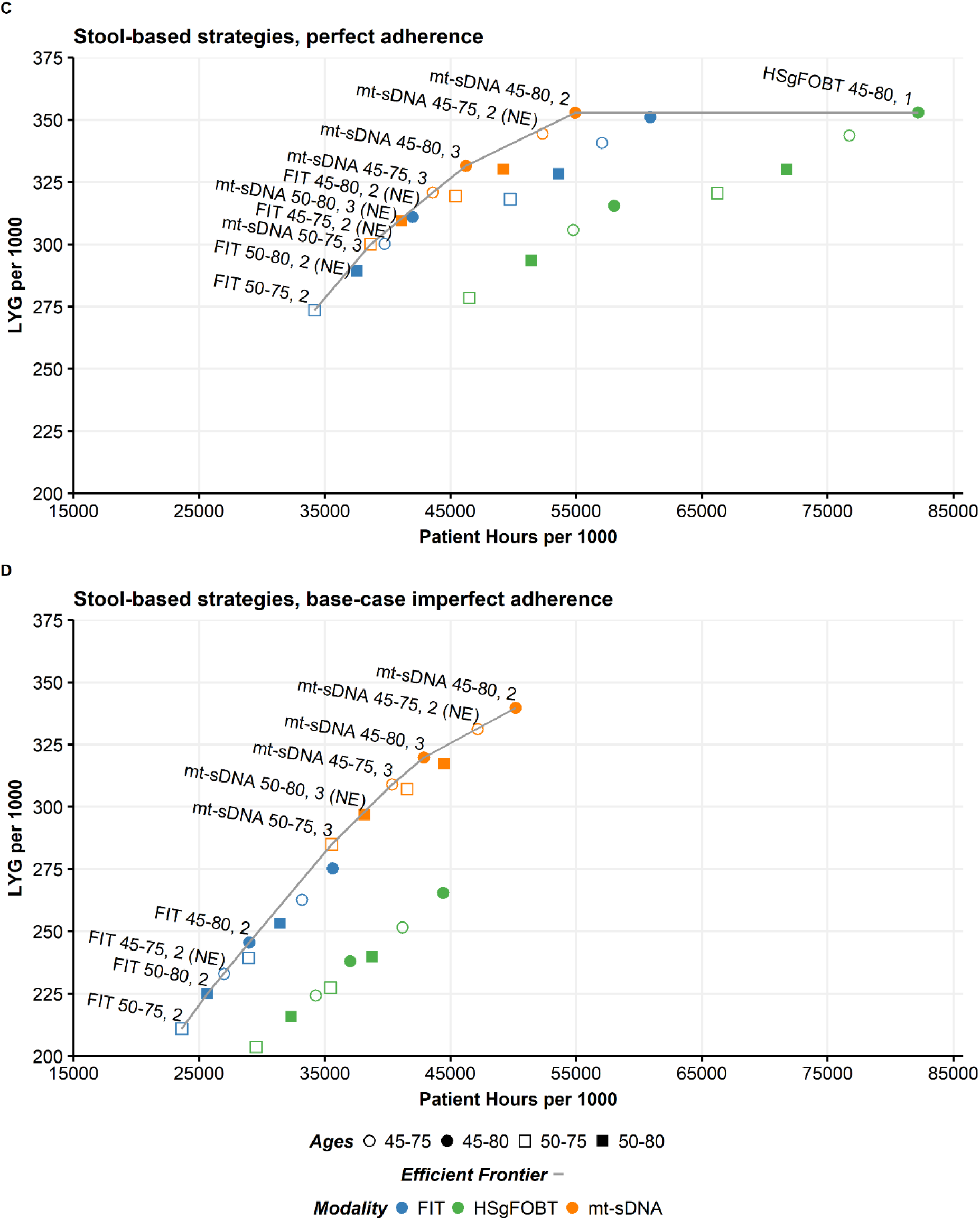
Life-years gained for individuals 40 years of age with stool-based tests by A) number of colonoscopies assuming perfect (100%) adherence or B) base-case imperfect adherence rates (40% FIT vs 34% HSgFOBT vs 70% mt-sDNA) and by C) patient hours related to the screening process assuming perfect (100%) adherence or D) base-case imperfect adherence (40% FIT vs 34% HSgFOBT vs 70% mt-sDNA). Results shown are per 1000 individuals free of diagnosed colorectal cancer at age 40 and screened starting at age 45 or 50 and ending at age 75 or 80 receiving biennial or triennial mt-sDNA, annual or biennial FIT, and annual or biennial HSgFOBT. NE, near-efficient.

The efficient frontiers by LYG relative to patient hours related to the screening process assuming perfect adherence or base-case imperfect adherence rates are shown in **Fig 3**. When shifting from perfect to base-case imperfect adherence rates, all but one of the strategies that are on or off the efficient frontier do not change (**Supplemental Tables S6 and S7**). Although there are no pre-established efficiency thresholds for this analysis, an absolute upper bound of an incremental 8,766 screening process hours per incremental LYG could be used, representing an equivalent tradeoff of burden (365.25 days/year x 24 hours/day=8,766 hours) and benefit, with other thresholds consisting of a fraction of this value. Assuming a screening window from 50–75, followed by recalculating efficiency comparisons, the annual HSgFOBT strategy on the frontier would also exceed the threshold with 16,807.5 incremental patient hours per incremental LYG compared with biennial mt-sDNA, which has an efficiency ratio of 350.0 hours per LYG and could be considered an optimal strategy (**Supplemental Table S8**). Similarly, at base-case imperfect adherence rates, mt-sDNA generally had preferable efficient ratios while offering the most LYG for a given screen window (**Supplemental Table S8**). There were no HSgFOBT strategies that were efficient or near-efficient. Similar results were obtained for a screening window from 45-75 (**Supplemental Table S8**).

### Analysis 3: Varying numbers of completed tests

Adherence analysis #3 assumed varying numbers of completed stool-based tests. Life-years gained per 1000 individuals screened between ages 50–75 was greater with up to 1, up to 5, or up to 9 mt-sDNA tests (68.2, 217.8, and 294.8, respectively) compared with equivalent numbers of FIT tests (41.4, 143.1, and 201.2; **Table 4**). An individual would have to take up to 21 FIT tests to reach approximately the same LYG as an individual who took up to 9 mt-sDNA tests (**Fig 4**). For up to 1, 5, or 9 tests, the incremental ratios of COL/LYG were favorable for mt-sDNA compared with FIT (**Table 4**) and are below the CISNET-accepted ratios of 39-65.

**Table 4.** Predicted outcomes per 1000 individuals screened from ages 50–75 compared with no screening.^a^

| Screening strategy | Randomly Assigned Number of Tests | Total Stool Tests | Total COLs | CRC Cases | CRC Deaths | LY with CRC | LYG | Incremental COL/ Incremental LYG vs FIT | Incidence Reduction | Mortality Reduction |
| --- | --- | --- | --- | --- | --- | --- | --- | --- | --- | --- |
| mt-sDNA, 50-75 | Up to 1 | 819 | 416 | 67.9 | 29.5 | 603.4 | 68.2 | 6.5 | 15.5% | 19.4% |
| FIT, 50-75 | Up to 1 | 813 | 243 | 73.7 | 32.5 | 631.9 | 41.4 |  | 8.2% | 11.3% |
| mt-sDNA, 50-75 | Up to 5 | 3,468 | 1,311 | 41.2 | 15.8 | 454.3 | 217.8 | 7.7 | 48.7% | 56.9% |
| FIT, 50-75 | Up to 5 | 3,710 | 735 | 55.8 | 22.3 | 564.8 | 143.1 |  | 30.6% | 39.1% |
| mt-sDNA, 50-75 | Up to 9 | 5,792 | 1,904 | 29.4 | 10.5 | 339.5 | 294.8 | 8.8 | 63.4% | 71.3% |
| FIT, 50-75 | Up to 9 | 6,250 | 1,084 | 45.2 | 17.1 | 502.9 | 201.2 |  | 43.8% | 53.3% |

COL, colonoscopy; CRC, colorectal cancer; FIT, fecal immunochemical test; LY, life-years; LYG, life-years gained; mt-sDNA, multitarget stool DNA test.
<sup>a</sup>Individuals were randomly assigned numbers of mt-sDNA (max=9 triennial tests during the screening window) or FIT (max=26 annual tests during the screening window).

**Fig 4.**
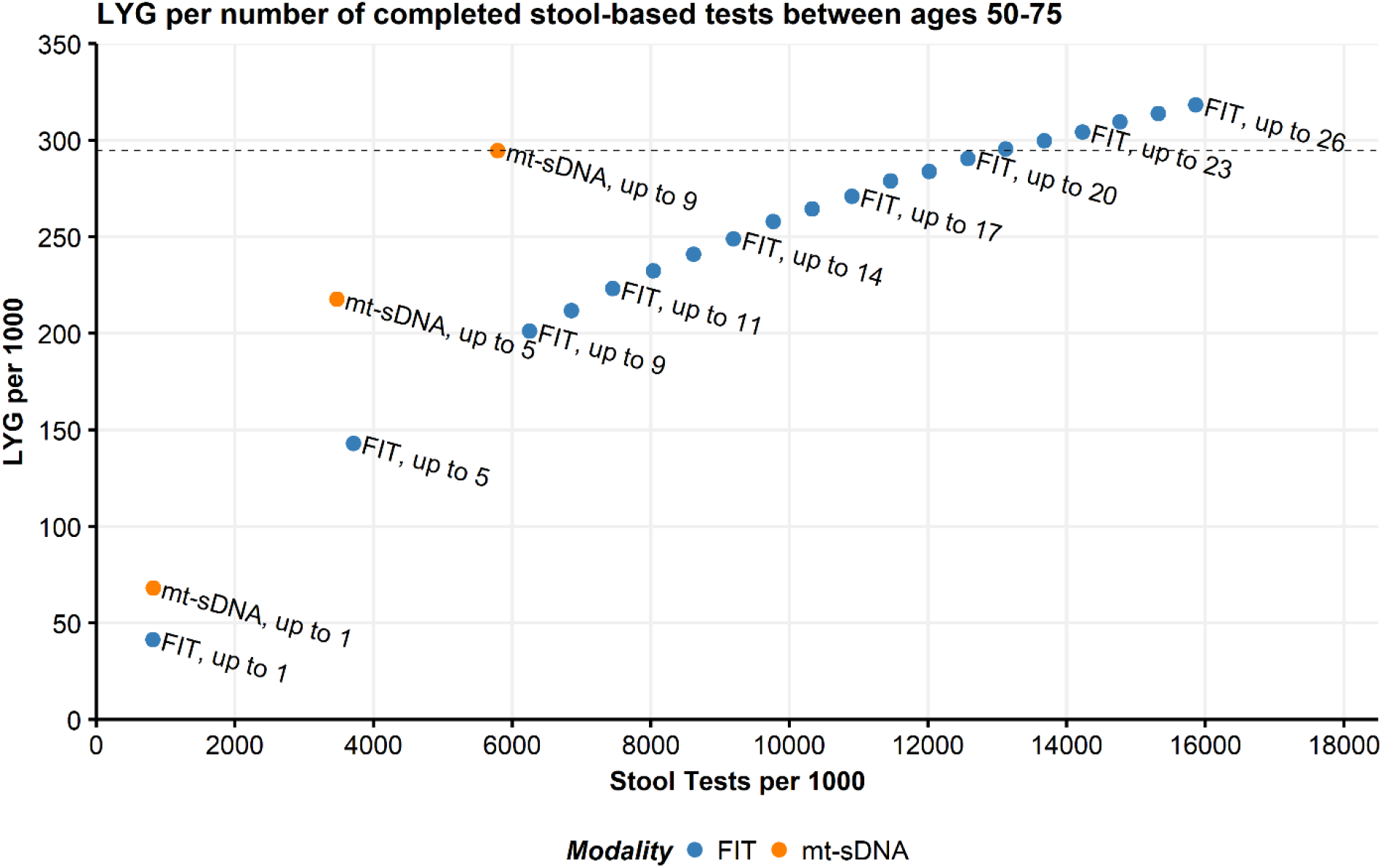
Predicted life-years gained (LYG) by total stool tests per 1000 individuals screened from ages 50–75 compared with no screening. Individuals were randomly assigned numbers of multitarget stool DNA (mt-sDNA; max=9 triennial tests during the screening window) or fecal immunochemical tests (FIT; max=26 annual tests during the screening window). The line indicates equivalent LYG with up to 9 mt-sDNA tests.

The reductions in CRC-related incidence and CRC-related mortality were also greater with up to 1, up to 5, or up to 9 mt-sDNA tests compared with equivalent numbers of FIT tests, whereas the number of total stool tests was similar between mt-sDNA and FIT when comparing equivalent numbers of tests (**Table 4**).

The pattern of results for analysis #3 in individuals screened between ages 45–75 was similar to those of individuals screened between ages 50–75 (**Supplemental Table S9**). An individual would have to take up to 25 FIT tests to reach approximately the same LYG as an individual who took up to 11 mt-sDNA tests (**Supplemental Fig S4**).

### Sensitivity analyses

#### Test performance sensitivity analyses

When granular adenoma size and location sensitivity values and age-based specificity values are used from DeeP-C (clinicaltrials.gov identifier, NCT01397747),[21] results from analysis #1 and #3 only modestly change. In analysis #1, when assuming perfect adherence in individuals screened between ages 50–75, LYG for triennial mt-sDNA increased from the primary analysis by 1.3% (303.8 vs 300.0) and total colonoscopies decreased by 4.3% (1872 vs 1957). LYG for annual FIT decreased by 1.0% (315.0 vs 318.1) and total colonoscopies decreased by 5.4% (1927 vs 2037) (**Supplemental Tables S2** and **S10**). At base-case imperfect adherence rates in individuals screened between ages 50–75, mt-sDNA resulted in a 19.6% increase in LYG (288.1) versus FIT (241.0; **Supplemental Fig S5**). In analysis #3, for up to 1, 5, or 9 tests, the incremental ratios of COL/LYG remained favorable for mt-sDNA compared with FIT and decreased by 4.6%, 6.5%, and 10.2%, respectively, from the primary analysis (**Table 4** and **Supplemental Table S11**). The number of FIT tests an individual would have to take to reach approximately the same LYG as an individual who took up to 9 mt-sDNA tests increased from 21 to 22 FIT tests (**Supplemental Fig S6**).

#### Efficiency frontier sensitivity analyses

For scenarios assuming 50% FIT vs 43% HSgFOBT vs 70% mt-sDNA adherence or 60% FIT vs 52% HSgFOBT vs 70% mt-sDNA adherence, the efficient frontiers showed only modest changes from the base-case imperfect adherence rate analysis. Both FIT and mt-sDNA continued to have efficient strategies while HSgFOBT did not (**Supplemental Fig S7**). When the screen window is fixed at 50-75 followed by recalculating efficiency comparisons, the recommendable strategy is biennial mt-sDNA with an efficiency ratio of 14.7 assuming 50% FIT vs 43% HSgFOBT vs 70% mt-sDNA adherence and 18.1 assuming 60% FIT vs 52% HSgFOBT vs 70% mt-sDNA adherence. Across the two sensitivity scenarios, only one strategy changed from (near)-efficient to not-efficient (triennial mt-sDNA, 50–75); this change occurred in the 60% FIT vs 52% HSgFOBT vs 70% mt-sDNA adherence scenario (**Supplemental Table S5**).

For the efficient frontiers by LYG relative to patient hours related to the screening process, the results were similar to those from the base-case imperfect adherence rate analysis and the strategies classified as either efficient or near-efficient were unchanged (**Supplemental Fig S7**). When the screen window is fixed followed by recalculating efficiency comparisons, conclusions regarding optimal strategy did not change (**Supplemental Table S8**).

Sensitivity analyses for analysis #2 using the CISNET fixed split adherence approach are shown in **Supplemental Fig S8**. The number of efficient strategies increases for mt-sDNA and decreases for FIT. HSgFOBT does not produce efficient strategies in either modeling approach. The CISNET fixed split adherence approach is more favorable to mt-sDNA strategies, as more of these strategies are on (or approach) the efficient frontier relative to the primary analysis (**Fig 3** and **Supplemental Fig S8**).

## Discussion

Using the CRC-AIM microsimulation model,[16, 17] a robust and comprehensive analysis was conducted to determine the impact of imperfect adherence on stool-based CRC screening outcomes. The results demonstrate that imperfect adherence has a substantial impact on the relative benefits and harms of guideline-endorsed, stool-based CRC screening strategies. The trade-offs between burden and benefit change when adherence rates are assumed to be less than 100%, shifting 4 of the 24 stool-based screening strategies from dominated to efficient or near-efficient (annual FIT 50-80; biennial and triennial mt-sDNA 45-75; triennial mt-sDNA 45-80). The model-recommended strategy also shifts from an annual FIT to a biennial mt-sDNA. At base-case imperfect adherence, mt-sDNA strategies are efficient/near-efficient, have efficiency ratios below accepted thresholds, and offer more LYG than FIT or HSgFOBT. When assuming patients are randomly adherent to an equal number of stool-based screening tests, more than twice the number of FIT tests are needed to match the equivalent amount of LYG as mt-sDNA. The LYG from FIT and HSgFOBT are more sensitive to changes in adherence assumptions than mt-sDNA because their lower neoplasia sensitivity requires more tests be completed for equivalent benefit.

Other modeling studies have observed substantive influence from imperfect CRC screening adherence on the simulated outcomes. Similar to the current analyses, another microsimulation model found that when reported adherence rather than perfect adherence was assumed, the LYG and reductions in CRC incidence and mortality were higher with triennial mt-sDNA compared with annual FIT or HSgFOBT.[10] In Zauber et al 2008[22], sensitivity analysis showed that when adherence was lowered from 100% to 50%, the LYG became higher for FIT and HSgFOBT compared with colonoscopy (mt-sDNA was not included in the analysis). Sensitivity analyses in Knudsen at al 2010[40] demonstrated that the optimal CRC screening modality can change when adherence is varied. Adherence to less-invasive tests appears to be an important, yet under-appreciated, factor when assessing the relative cost-effectiveness of CRC screening.[42]

In routine clinical practice, adherence to test-specific CRC screening strategies is imperfect and varies across patient subgroups.[12–15] Even for a single individual, the likelihood to adhere may change over time because of a variety of factors (e.g., education about screening, life events, etc.). Furthermore, patients may switch between different screening modalities and adherence may decline over time. Although imperfect adherence in real-world settings is recognized,[4] perfect adherence is usually used in CRC screening microsimulations because there is no universally accepted methodology to model imperfect adherence and all of the methods used have challenges. One method often utilized by CISNET is to assume a proportion of a population is perfectly adherent to a test, and the remaining proportion are never adherent (fixed split adherence approach).[39–41] This effectively results in a weighted average of outcomes for 100% adherence and natural history (i.e., never-adherent).[40] Commonly used values for this imperfect adherence assumption generally range between 50%-60%, loosely corresponding to the proportion of individuals up-to-date with CRC screening.[39–41] A challenge of this approach is that the proportion of individuals up-to-date for a CRC test does not necessarily correspond to constant, perfect longitudinal adherence, it only corresponds to a proportion of individuals who have previously undergone a screen at some point within a screen modality’s testing interval, ignoring whether the screen was taken on time (e.g. at age 60, the proportion who took a colonoscopy at age 50, rather than the proportion that took the test over the last 10 years). Additionally, evidence suggests that it is incorrect to assume that individuals who previously took a screening test will consistently take it in future years.[43, 44] The fixed split adherence approach also assumes that individuals who are not adherent will never take a screening test, which is also unrealistic.[34]

The current analysis models imperfect adherence for stool tests using cross-sectional, first-round, participation rates. This approach assumes that if an individual does not adhere to a test, that the individual has the same fixed probability to adhere the following year and that for individuals that do adhere and the test is negative, the individual has the same fixed annual likelihood to adhere in future screen opportunities. Over infinite time, there are no individuals who never or perfectly adhere. Many more individuals get screened using this approach compared with the fixed split adherence approach.

The most realistic simulation of adherence may be somewhere in between the two methods. One approach has been to model adherence as a mixed distribution of individuals who never adhere, perfectly adhere, and imperfectly adhere, with the imperfectly adherent group further sub-stratified into different levels of adherence likelihood.[22, 45] Given a lack of longitudinal adherence data to act as calibration and validation targets, the inter and intra-distributions of these groups are difficult to quantify, especially if differential adherence is assumed across strategies.

Despite the challenges in modeling real-world adherence, the strength of this analysis is that multiple modeling approaches were conducted and many sensitivities were explored to address the challenges. For analysis #1, a spectrum of adherence in 10% intervals was applied. Although this approach does not reflect a real-world population, it is a simplified and thorough way to show the impact of imperfect adherence. For analysis #2, the goal was to apply real-world first-round differential adherence rates. Therefore, we conducted a critical assessment of meta-analyses and retrospective cross-sectional data, which, although limited by some caveats, indicated that actual adherence for annual FIT is likely ~40% based on a weighted average (accounting for heterogeneity) of 42% adherence from studies included in a meta-analysis[29]. This rate is similar to that determined from a large database of Veteran’s Affairs data in the US (42.6%).[46] Based on meta-analysis data, annual HSgFOBT is ~16%[29] to 21%[38] relatively lower than FIT, thus, HSgFOBT adherence rates of 16% lower relative to FIT were assumed. The 70% adherence rate with triennial mt-sDNA was determined based on a large (n=368,494) retrospective analysis of Medicare beneficiaries from diverse geographic regions.[30]

The higher adherence rate with mt-sDNA compared with FIT and HSgFOBT is likely related to multiple factors. Patient navigation programs are always available for patients using mt-sDNA[30] and such programs have been shown to increase adherence.[47–50] Differences in the practical aspects of completing the stool-based tests may also play a role. For example, HSgFOBT requires diet modification and samples from multiple bowel movements, unlike FIT and mt-sDNA.[51] Although the retrospective analysis used to determine the mt-sDNA adherence was conducted in the Medicare population and older patients are known to be more adherent to CRC screening,[12, 52] within the analyzed population there were no age-related trends for mt-sDNA adherence nor were any trends observed based on the type of Medicare coverage.[30]

A sensitivity analysis of analysis #2 was conducted using adherence rates of 50% and 60% for FIT (and corresponding 16% relatively lower rates for HSgFOBT). Adherence values of 50% and 60% were based on a weighted average (accounting for heterogeneity) of 56% adherence for first-round participation rates reviewed by the US Multi-Society Task Force on Colorectal Cancer.[33] Adherence of approximately 50% has been observed by Kaiser Permamente in a retrospective cohort study of 323,349 members.[14] The Kaiser Permanente analysis was within the context of an organized screening program that included patient navigation.[14] Such programs are not universally available to FIT patients in the US. The highest reported rates of FIT adherence were reported in Dutch studies (~60%).[34–37] Some of the Dutch studies were in the context of a CRC screening program and may not be reflective of a US population. Socioeconomic, cultural, and geographic factors (i.e., patients were selected only from municipalities), as well as national policies may have contributed to high adherence rates.[53] Results from the sensitivity analyses using higher FIT and HSgFOBT adherence rates are in agreement with the base-case imperfect adherence analysis in that biennial mt-sDNA is the model recommendable strategy within a 50-75 year screening window.

“Screening fatigue”, defined as missing a test after receiving several negative CRC tests, is a potential problem that reduces the effectiveness of screening.[54] A CRC model estimated that screening fatigue could considerably reduce screening effectiveness.[55] In analysis #3, individuals are only randomly adherent to a fixed number of completed stool tests, similar to a patient with screening fatigue who is only willing to take a limited number of stool tests. In this context, a patient would need to be willing to take approximately 21 FIT tests to produce the same LYG as 9 mt-sDNA tests. A retrospective analysis of individuals with commercial insurance or Medicare in the US found that only 268/17,174 (1.56%) of the individuals who received any annual FIT/HSgFOBT test (with no sigmoidoscopy) received at least 1 annual FIT/FOBT test per year over 10 years.[56] This excludes individuals who had a follow-up colonoscopy from a false-positive. Therefore, it is unlikely that many patients would complete 21 negative FIT tests in a 25-year period.

Efficient frontiers are a way to evaluate the trade-offs between the benefits and harms from screening tests. Although the CISNET modeling group commonly uses total colonoscopies for efficient frontier analysis, they have also recognized the need to assess alternate metrics for patient burden when recommending modeling strategies.[4] The current analysis evaluated efficient frontiers not only from a healthcare burden perspective (e.g., LYG relative to number of colonoscopies), but also from a patient burden perspective (e.g., LYG relative to number of patient hours related to the screening process). At base-case imperfect adherence rates, mt-sDNA generally had preferable efficient ratios while offering the most LYG when compared with FIT or HSgFOBT from both the healthcare and patient burden perspectives. Looking at other measures of burden may be complementary to using number of colonoscopies to provide an additional perspective for efficiency frontiers.

The CRC-AIM model is limited in that, like the CISNET models, it does not account for serrated polyps, the natural history of which differs from the typical adenoma-carcinoma sequence, and may account for up to 30% of CRC.[57, 58] Sessile serrated polyps are detected with good sensitivity by mt-sDNA but not FIT.[59, 60] Although not explicitly modeled, hyperplastic polyp detection is reflected in the false-positive rate (1-specificity) for colonoscopy. The model also does not account for performance of colonoscopy (adenoma sensitivity) in the real-world that varies by endoscopist[61] and by screening versus follow-up.[62] Finally, to be comparable to CISNET models, the primary CRC-AIM analysis was limited to using CISNET test characteristic parameters. In sensitivity analyses of analysis #1 and #3, inputs for sensitivity by adenoma size/location and specificity by age for mt-sDNA and FIT were used after obtaining clinical data from the sponsor of the study published by Imperiale et al (DeeP-C; clinicaltrials.gov identifier, NCT01397747).[21] Although the changes to the predicted outcomes were relatively small, more granular test performance is also needed for other screening modalities for equivalent comparisons because small changes may impact efficiency frontiers.

## Conclusion

Based on the CRC-AIM model, all stool-based screening strategies decrease CRC-related incidence and mortality compared with no screening, regardless of adherence assumptions. Conclusions about the comparative tradeoffs between burden and benefit that are used to inform CRC screening guidelines change when adherence rates are assumed to be less than 100%. At imperfect adherence rates, mt-sDNA provides more LYG than FIT or HSgFOBT at an acceptable tradeoff in screening burden. LYG from FIT and HSgFOBT are more sensitive to changes in adherence assumptions than mt-sDNA because they require more tests be completed for equivalent benefit.

Real-world adherence to stool-based testing is imperfect and should be simulated using realistic inputs in comparative effectiveness modeling to more accurately assess the benefits of screening. Physicians and guidelines should consider the impact of imperfect adherence when making recommendations to patients.

## Acknowledgements

Medical writing and editorial assistance were provided by Erin P. Scott, PhD, of Maple Health Group, LLC, funded by Exact Sciences Corporation.

## Supporting Information Captions

### Supporting Methods

**Table S1. Screening characteristics for Deep-C (clinicaltrials.gov identifier, NCT01397747) sensitivity analyses using granular adenoma size**.

**Table S2. Screening outcomes per 1000 individuals by adherence rate for triennial mt-sDNA, annual FIT, and annual HSgFOBT in individuals free of diagnosed colorectal cancer at age 40 and screened between ages 50–75 years or 45–75 years**.

**Table S3. Outcomes and efficiency ratios of LYG relative to number of colonoscopies assuming perfect (100%) adherence**. Results are ordered by total colonoscopies. Results shown are per 1000 individuals free of diagnosed colorectal cancer at age 40 and screened starting at age 45 or 50 and ending at age 75 or 80 receiving biennial or triennial mt-sDNA, annual or biennial FIT, and annual or biennial HSgFOBT. Bold row is the model-recommended strategy.

**Table S4. Outcomes and efficiency ratios of LYG relative to number of colonoscopies at base-case imperfect adherence rates of 40% FIT vs 34% HSgFOBT vs 70% mt-sDNA**. Results are ordered by total colonoscopies. Results shown are per 1000 individuals free of diagnosed colorectal cancer at age 40 and screened starting at age 45 or 50 and ending at age 75 or 80 receiving biennial or triennial mt-sDNA, annual or biennial FIT, and annual or biennial HSgFOBT. Gray highlight indicates shift from dominated to efficient or near-efficient from 100% adherence assumption. Italics indicates shift from efficient or near-efficient to dominated from 100% adherence assumption. Bold row is the model-recommended strategy.

**Table S5. Incremental efficiency ratios for number of colonoscopies at a fixed screening window of 50–75 or 45–75 assuming perfect (100%) adherence, base-case imperfect adherence rates of 40% FIT vs 34% HSgFOBT vs 70% mt-sDNA, adherence of 50% FIT vs 43% HSgFOBT vs 70% mt-sDNA, or adherence of 60% FIT vs 52% HSgFOBT vs 70% mt-sDNA**. Results are ordered by total colonoscopies. Results shown are per 1000 individuals free of diagnosed colorectal cancer receiving biennial or triennial mt-sDNA, annual or biennial FIT, and annual or biennial HSgFOBT.

**Table S6. Outcomes and efficiency ratios based on patient hours related to the screening process assuming perfect (100%) adherence**. Results are ordered by patient hours. Results shown are per 1000 individuals free of diagnosed colorectal cancer at age 40 and screened starting at age 45 or 50 and ending at age 75 or 80 receiving biennial or triennial mt-sDNA, annual or biennial FIT, and annual or biennial HSgFOBT. Bold row is the model-recommended strategy.

**Table S7. Outcomes and efficiency ratios based on patient hours related to the screening process at base-case imperfect adherence rates of 40% FIT vs 34% HSgFOBT vs 70% mt-sDNA**. Results are ordered by patient hours. Results shown are per 1000 individuals free of diagnosed colorectal cancer at age 40 and screened starting at age 45 or 50 and ending at age 75 or 80 receiving biennial or triennial mt-sDNA, annual or biennial FIT, and annual or biennial HSgFOBT. Italics indicates shift from efficient or near-efficient to dominated from 100% adherence assumption. Bold row is the model-recommended strategy.

**Table S8. Incremental efficiency ratios for patient hours related to the screening process at a fixed screening window of 50–75 or 45–75 assuming perfect (100%) adherence, base-case imperfect adherence rates of 40% FIT vs 34% HSgFOBT vs 70% mt-sDNA, 50% FIT vs 43% HSgFOBT vs 70% mt-sDNA adherence, or 60% FIT vs 52% HSgFOBT vs 70% mt-sDNA adherence**. Results are ordered by patient hours. Results shown are per 1000 individuals free of diagnosed colorectal cancer receiving biennial or triennial mt-sDNA, annual or biennial FIT, and annual or biennial HSgFOBT.

**Table S9. Predicted outcomes per 1000 individuals screened from ages 45–75 compared with no screening**. Individuals were randomly assigned numbers of mt-sDNA (max=11 triennial tests during the screening window) or FIT (max=31 annual tests during the screening window).

**Table S10. Deep-C (clinicaltrials.gov identifier, NCT01397747) sensitivity analysis for screening outcomes per 1000 individuals by adherence rate for triennial mt-sDNA and annual FIT in individuals free of diagnosed colorectal cancer at age 40 and screened between ages 50–75 years or 45-–75 years**.

**Table S11. Deep-C (clinicaltrials.gov identifier, NCT01397747) sensitivity analysis of predicted outcomes per 1000 individuals screened from ages 50–75 compared with no screening**. Individuals were randomly assigned numbers of mt-sDNA (max=9 triennial tests during the screening window) or FIT (max=26 annual tests during the screening window).

**Figure S1. Age-specific risks of complications from colonoscopy with polypectomy used in the analysis**. Complications include serious gastrointestinal events, other gastrointestinal events, and cardiovascular events. We assume that the only harms from screening arise from a colonoscopy with polypectomy, whether it be for screening, follow-up, or surveillance, or for the diagnosis of a symptomatic cancer. We assume no risk of harms from stool-based tests, nor from bowel preparation.[28] The risks of complications from colonoscopy are from an analysis by van Hees et al.,[26] which was an extension of an analysis by Warren et al.[27] In those studies, colonoscopy without polypectomy was not associated with an excess risk of complications, relative to a matched control group that did not have colonoscopy. Reproduced and adapted with permission from Knudsen et al, 2016.[4]

**Figure S2. Percent difference in predicted life-years gained (LYG) per 1000 individuals by adherence rate for triennial multitarget stool DNA (mt-sDNA) versus annual fecal immunochemical test (FIT) in individuals free of diagnosed colorectal cancer at age 40 and screened between ages 45–75 years**. White boxes indicate <10% difference between tests. Light gray boxes indicate ≥10% positive difference with mt-sDNA versus FIT. Dark gray boxes indicate ≥10% negative difference with mt-sDNA versus FIT. Outlined box indicates base-case imperfect adherence rates.

**Figure S3. A) Predicted life-years gained (LYG), B) CRC-related incidence and mortality reduction, and C) total stool tests and colonoscopies (COL) per 1000 individuals screened from ages 45–75 compared with no screening assuming perfect (100%) adherence rates to annual FIT, annual HSgFOBT, and triennial mt-sDNA or base-case imperfect adherence (40% FIT vs 34% HSgFOBT vs 70% mt-sDNA**).

**Figure S4. Predicted life-years gained (LYG) by total stool tests per 1000 individuals screened from ages 45–75 compared with no screening**. Individuals were randomly assigned numbers of multitarget stool DNA (mt-sDNA; max=11 triennial tests during the screening window) or fecal immunochemical tests (FIT; max=31 annual tests during the screening window). The line indicates equivalent LYG with up to 11 mt-sDNA tests.

**Figure S5. Deep-C (clinicaltrials.gov identifier, NCT01397747) sensitivity analysis of percent difference in predicted life-years gained (LYG) per 1000 individuals by adherence rate for triennial multitarget stool DNA (mt-sDNA) versus annual fecal immunochemical test (FIT) in individuals free of diagnosed colorectal cancer at age 40 and screened between ages 50–75 years**. White boxes indicate <10% difference between tests. Light gray boxes indicate ≥10% positive difference with mt-sDNA versus FIT. Dark gray boxes indicate ≥10% negative difference with mt-sDNA versus FIT. Outlined box indicates base-case imperfect adherence rates.

**Figure S6. Deep-C (clinicaltrials.gov identifier, NCT01397747) sensitivity analysis of predicted life-years gained (LYG) by total stool tests per 1000 individuals screened from ages 50–75 compared with no screening**. Individuals were randomly assigned numbers of multitarget stool DNA (mt-sDNA; max=9 triennial tests during the screening window) or fecal immunochemical tests (FIT; max=26 annual tests during the screening window). The line indicates equivalent LYG with up to 9 mt-sDNA tests.

**Figure S7. Sensitivity analysis of life-years gained for individuals 40 years of age with stool-based tests A) number of colonoscopies assuming 50% FIT vs 43% HSgFOBT vs 70% mt-sDNA adherence or B) assuming 60% FIT vs 52% HSgFOBT vs 70% mt-sDNA adherence and by C) patient hours related to the screening process and assuming 50% FIT vs 43% HSgFOBT vs 70% mt-sDNA adherence or D) assuming 60% FIT vs 52% HSgFOBT vs 70% mt-sDNA adherence**. Results shown are per 1000 individuals free of diagnosed colorectal cancer at age 40 and screened starting at age 45 or 50 and ending at age 75 or 80 receiving biennial or triennial mt-sDNA, annual or biennial FIT, and annual or biennial HSgFOBT. NE, near-efficient.

**Figure S8. Life-years gained for individuals 40 years of age with stool-based tests by A) number of colonoscopies or B) patient hours related to the screening process assuming base-case imperfect adherence rates of 40% FIT vs 34% HSgFOBT vs 70% mt-sDNA using the CISNET fixed split adherence approach**. Results shown are per 1000 individuals free of diagnosed colorectal cancer at age 40 and screened starting at age 45 or 50 and ending at age 75 or 80 receiving biennial or triennial mt-sDNA, annual or biennial FIT, and annual or biennial HSgFOBT. NE, near efficient.

## Notes

**Funding:** Financial support for this study was provided by a contract with Exact Sciences Corporation. https://www.exactsciences.com/ The funding agreement ensured the authors’ independence in designing the study, interpreting the data, writing, and publishing the report. LS, MM, ABO, and MP are employees of the study sponsor and received a salary.

**Competing interests:** The study was not related to any patents, patent applications, or products in development.

https://github.com/CRCAIM/CRC-AIM-Public

### Competing Interest Statement

A. Piscitello is an employee of EmpiriQA, LLC, which provides consulting services to Exact Sciences Corporation. L. Saoud, M. Matney, A.B. Ozbay, and M. Parton are employees of Exact Sciences Corporation. A.M. Fendrick has been a consultant for AbbVie, Amgen, Centivo, Community Oncology Association, Covered California, EmblemHealth, Exact Sciences, Freedman Health, GRAIL, Harvard University, Health & Wellness Innovations, Health at Scale Technologies, MedZed, Penguin Pay, Risalto, Sempre Health, the State of Minnesota, U.S. Department of Defense, Virginia Center for Health Innovation, Wellth, and Zansors; has received research support from the Agency for Healthcare Research and Quality, Gary and Mary West Health Policy Center, Arnold Ventures, National Pharmaceutical Council, Patient-Centered Outcomes Research Institute, Pharmaceutical Research and Manufacturers of America, the Robert Wood Johnson Foundation, the State of Michigan, and the Centers for Medicare and Medicaid Services. B.J. Borah has nothing to disclose. K. Hassmiller Lich has nothing to disclose. P.J. Limburg serves as Chief Medical Officer for Screening at Exact Sciences through a contracted services agreement with Mayo Clinic. Dr. Limburg and Mayo Clinic have contractual rights to receive royalties through this agreement.

