## Supplemental information for "Estimating the impact of differential adherence on the comparative effectiveness of stool-based colorectal cancer screening using the CRC-AIM microsimulation model"

### Supplemental Material

#### Methods

*Microsimulation model*

Adenomas within an individual are generated using a non-homogenous Poisson process based on sex, age, and person-specific risk and assigned a location in either the colon or rectum.[1, 2] Adenomas growth rates based on location are assumed to follow a non-linear growth curve. The cumulative probability of an adenoma transitioning to preclinical CRC is assumed to be a function of adenoma size, age at adenoma initiation, sex, and location of the adenoma. After the adenoma transitions to preclinical CRC, the preclinical CRC is assigned an initial size of 0.5 mm, the sojourn time is determined, and the size of the preclinical CRC upon reaching sojourn time is sampled from a Surveillance, Epidemiology, and End Results (SEER) distribution of CRC sizes. The model assumes simple exponential growth of the CRC. Cancer stage at diagnosis and detection is determined based on CRC size, and the stage of CRC at detection is further determined using a multinomial logistic regression model.[1, 2] Sex-specific cohort life tables from the years 1900-2010 are used for the all-cause (non-CRC) mortality rates. CRC stage-specific survival is based on parametric regression models developed from SEER data for cancers diagnosed from 2000-2003.

For the CRC-AIM screening component it is assumed that CRC screening facilitates the detection and removal of adenomas and preclinical lesions and that the ability of a lesion to be detected is dependent on the sensitivity and reach of the screening test.[3, 4] Screening frequency and adherence rates, as well as the sensitivity and specificity of the screening test, impact the effectiveness of the test. The sensitivity inputs for stool-based tests are per person and are based on the characteristics of the most advanced lesion. The sensitivity inputs for structural tests are per lesion and potential detection of lesions depend on the reach of the test. False positives can occur. Complications (e.g., serious and non-serious gastrointestinal events, cardiovascular events) due to polypectomies can arise and are part of the screening component.

The model assumes that in all cases a follow-up colonoscopy occurs after any positive non-colonoscopy screening test.[3, 4] After a negative follow-up colonoscopy, individuals return to their original non-colonoscopy screening test and the next screening is due in 10 years. After a positive follow-up colonoscopy, individuals enter a surveillance colonoscopy period where the next colonoscopy is based on the findings of the latest colonoscopy and continues until at least age 85. Individuals with preclinical lesions that become symptomatic based on sojourn time expiration receive a diagnostic colonoscopy.

#### Table S1. Screening characteristics for Deep-C (clinicaltrials.gov identifier, NCT01397747) sensitivity analyses using granular adenoma size.

| | Sensitivity | | | | Specificity | | | | |
|---|---|---|---|---|---|---|---|---|---|
| | Adenomas, mm | | | Cancer Stage | Age, y | | | | |
| | <6 | 6–9 | ≥10 | I–IV | <60 | 60–64 | 65–69 | 70–74 | 75+ |
| FIT, % (n/N) | 6.0% (117/1944) | 11.1% (113/1015) | 24.6% (170/691) | 73.8% (48/65) | 97.8% (1361/1392) | 96.5% (355/368) | 95.6% (1483/1551) | 96.7% (697/721) | 93.9% (399/425) |
| 95% CI | 5.0% – 7.2% | 9.3% – 13.2% | 21.4% – 28.0% | 61.5% – 84.0% | 96.9% – 98.5% | 94.0% – 98.1% | 94.5% – 96.6% | 95.1% – 97.9% | 91.2% – 96.0% |
| mt-sDNA, % (n/N) | 14.9% (290/1944) | 22.5% (228/1015) | 43.6% (301/691) | 92.3% (60/65) | 94.4% (1314/1392) | 92.4% (340/368) | 89.1% (1382/1551) | 86.1% (621/721) | 81.2% (345/425) |
| 95% CI | 13.4% – 16.6% | 19.9% – 25.2% | 39.8% – 47.4% | 83.0% – 97.5% | 93.1% – 95.5% | 89.2% – 94.9% | 87.4% – 90.6% | 83.4% – 88.6% | 77.1% – 84.8% |

FIT, fecal immunochemical test; mt-sDNA, multitarget stool DNA test.

2-sided 95% confidence intervals calculated using the Exact (Clopper-Pearson) method.

**Table S2. Screening outcomes per 1000 individuals by adherence rate for triennial mt-sDNA, annual FIT, and annual HSgFOBT in individuals free of diagnosed colorectal cancer at age 40 and screened between ages 50–75 years or 45–75 years.**

| Screening Strategy and Adherence Rate | Stool Tests | Follow-up COLs | Surveillance COLs | Total COLs | CRC Cases | CRC Deaths | LY with CRC | LYG | Incidence Reduction | Mortality Reduction |
|---|---|---|---|---|---|---|---|---|---|---|
| No screening | 0 | 0 | 0 | 80 | 80.3 | 36.6 | 646.0 | 0.0 | 0.0% | 0.0% |
| mt-sDNA 50-75, 3 | | | | | | | | | | |
| 10% | 1,782 | 268 | 452 | 770 | 57.6 | 24.6 | 526.9 | 133.1 | 28.3% | 33.0% |
| 20% | 2,925 | 424 | 684 | 1,145 | 46.8 | 18.9 | 464.0 | 196.0 | 41.7% | 48.3% |
| 30% | 3,717 | 527 | 825 | 1,382 | 40.6 | 15.9 | 422.0 | 231.1 | 49.4% | 56.5% |
| 40% | 4,304 | 601 | 915 | 1,541 | 36.9 | 14.0 | 397.0 | 253.8 | 54.1% | 61.7% |
| 50% | 4,751 | 657 | 978 | 1,659 | 34.4 | 12.8 | 378.5 | 268.1 | 57.2% | 65.0% |
| 60% | 5,107 | 701 | 1,024 | 1,747 | 32.6 | 12.0 | 364.0 | 278.3 | 59.4% | 67.2% |
| 70% | 5,397 | 734 | 1,059 | 1,813 | 31.3 | 11.4 | 353.4 | 285.1 | 61.1% | 68.8% |
| 80% | 5,641 | 763 | 1,085 | 1,867 | 30.4 | 11.0 | 346.9 | 290.2 | 62.2% | 70.0% |
| 90% | 5,864 | 789 | 1,108 | 1,916 | 29.4 | 10.6 | 339.5 | 295.3 | 63.4% | 71.2% |
| 100% | 6,076 | 814 | 1,126 | 1,957 | 28.6 | 10.2 | 333.0 | 300.0 | 64.4% | 72.2% |
| FIT 50-75, 1 | | | | | | | | | | |
| 10% | 2,174 | 142 | 285 | 486 | 65.5 | 28.1 | 582.6 | 96.3 | 18.5% | 23.2% |
| 20% | 4,162 | 258 | 500 | 802 | 55.1 | 22.4 | 530.7 | 160.2 | 31.5% | 38.8% |
| 30% | 5,980 | 356 | 666 | 1,057 | 47.2 | 18.4 | 483.8 | 206.5 | 41.3% | 49.8% |
| 40% | 7,672 | 441 | 799 | 1,269 | 41.3 | 15.5 | 444.4 | 239.4 | 48.6% | 57.6% |
| 50% | 9,246 | 518 | 903 | 1,444 | 36.9 | 13.5 | 411.5 | 263.0 | 54.1% | 63.1% |
| 60% | 10,727 | 585 | 987 | 1,594 | 33.6 | 12.0 | 384.9 | 279.5 | 58.2% | 67.2% |
| 70% | 12,120 | 648 | 1,060 | 1,726 | 30.8 | 10.9 | 360.5 | 293.0 | 61.6% | 70.3% |
| 80% | 13,432 | 705 | 1,120 | 1,842 | 28.7 | 9.9 | 342.8 | 304.4 | 64.3% | 72.9% |
| 90% | 14,673 | 759 | 1,170 | 1,945 | 26.9 | 9.3 | 323.7 | 310.9 | 66.5% | 74.6% |
| 100% | 15,856 | 808 | 1,214 | 2,037 | 25.4 | 8.7 | 308.9 | 318.1 | 68.4% | 76.3% |
| HSgFOBT 50-75, 1 | | | | | | | | | | |
| 10% | 2,127 | 208 | 306 | 572 | 64.4 | 27.7 | 572.4 | 99.8 | 19.9% | 24.4% |
| 20% | 3,981 | 379 | 532 | 954 | 53.1 | 21.7 | 511.5 | 166.3 | 33.8% | 40.6% |
| 30% | 5,612 | 520 | 701 | 1,255 | 45.3 | 17.8 | 463.8 | 211.9 | 43.7% | 51.5% |
| 40% | 7,074 | 642 | 833 | 1,503 | 39.3 | 14.9 | 422.0 | 244.2 | 51.1% | 59.3% |
| 50% | 8,373 | 752 | 934 | 1,709 | 34.9 | 12.9 | 390.0 | 268.0 | 56.5% | 64.9% |
| 60% | 9,558 | 846 | 1,016 | 1,883 | 31.7 | 11.5 | 361.5 | 284.9 | 60.6% | 68.7% |
| 70% | 10,631 | 933 | 1,084 | 2,034 | 29.1 | 10.4 | 338.2 | 296.6 | 63.7% | 71.6% |
| 80% | 11,609 | 1,011 | 1,139 | 2,167 | 27.1 | 9.5 | 321.6 | 307.3 | 66.3% | 74.0% |
| 90% | 12,505 | 1,081 | 1,188 | 2,285 | 25.4 | 8.9 | 302.8 | 314.2 | 68.4% | 75.7% |
| 100% | 13,337 | 1,146 | 1,227 | 2,388 | 24.0 | 8.4 | 288.4 | 320.6 | 70.1% | 77.0% |
| mt-sDNA 45-75, 3 | | | | | | | | | | |
| 10% | 2,155 | 309 | 523 | 880 | 55.0 | 23.3 | 499.9 | 151.1 | 31.5% | 36.3% |
| 20% | 3,522 | 491 | 781 | 1,306 | 43.6 | 17.5 | 426.7 | 220.4 | 45.8% | 52.2% |
| 30% | 4,472 | 609 | 929 | 1,565 | 37.4 | 14.6 | 381.5 | 255.7 | 53.4% | 60.3% |
| 40% | 5,171 | 696 | 1,022 | 1,742 | 33.9 | 12.8 | 355.2 | 277.5 | 57.8% | 65.1% |
| 50% | 5,711 | 759 | 1,089 | 1,869 | 31.4 | 11.7 | 334.6 | 291.1 | 60.9% | 68.1% |

| | | | | | | | | | | |
|---|---|---|---|---|---|---|---|---|---|---|
| 60% | 6,138 | 809 | 1,138 | 1,967 | 29.7 | 10.9 | 318.8 | 301.3 | 63.1% | 70.3% |
| 70% | 6,485 | 851 | 1,174 | 2,043 | 28.4 | 10.3 | 308.8 | 309.0 | 64.6% | 71.9% |
| 80% | 6,771 | 883 | 1,202 | 2,103 | 27.5 | 9.9 | 298.9 | 312.7 | 65.8% | 72.9% |
| 90% | 6,995 | 911 | 1,224 | 2,151 | 26.8 | 9.6 | 292.6 | 317.0 | 66.7% | 73.8% |
| 100% | 7,213 | 935 | 1,245 | 2,196 | 26.1 | 9.3 | 287.4 | 320.9 | 67.5% | 74.6% |
| FIT 45-75, 1 | | | | | | | | | | |
| 10% | 2,637 | 162 | 331 | 550 | 63.7 | 27.3 | 565.1 | 110.2 | 20.7% | 25.5% |
| 20% | 5,035 | 297 | 574 | 913 | 52.3 | 21.2 | 501.6 | 180.4 | 34.8% | 42.0% |
| 30% | 7,239 | 408 | 757 | 1,198 | 44.2 | 17.1 | 448.1 | 229.9 | 45.0% | 53.3% |
| 40% | 9,293 | 507 | 900 | 1,433 | 38.2 | 14.3 | 402.9 | 262.7 | 52.4% | 60.9% |
| 50% | 11,191 | 595 | 1,011 | 1,628 | 33.9 | 12.3 | 368.4 | 286.5 | 57.8% | 66.4% |
| 60% | 12,978 | 675 | 1,101 | 1,795 | 30.5 | 10.8 | 338.9 | 304.3 | 62.0% | 70.5% |
| 70% | 14,663 | 747 | 1,177 | 1,941 | 27.9 | 9.7 | 314.6 | 317.6 | 65.3% | 73.5% |
| 80% | 16,249 | 814 | 1,238 | 2,067 | 25.8 | 8.9 | 293.6 | 327.6 | 67.9% | 75.8% |
| 90% | 17,751 | 877 | 1,292 | 2,183 | 24.1 | 8.2 | 274.5 | 334.7 | 70.0% | 77.5% |
| 100% | 19,182 | 935 | 1,336 | 2,285 | 22.8 | 7.7 | 261.3 | 340.7 | 71.7% | 78.9% |
| HSgFOBT 45-75, 1 | | | | | | | | | | |
| 10% | 2,575 | 244 | 355 | 654 | 62.4 | 26.8 | 553.2 | 114.7 | 22.3% | 26.8% |
| 20% | 4,808 | 445 | 608 | 1,093 | 50.3 | 20.5 | 480.2 | 186.6 | 37.3% | 44.0% |
| 30% | 6,772 | 609 | 794 | 1,434 | 42.1 | 16.4 | 423.2 | 235.3 | 47.6% | 55.2% |
| 40% | 8,520 | 756 | 934 | 1,715 | 36.1 | 13.6 | 378.8 | 270.1 | 55.1% | 62.9% |
| 50% | 10,086 | 881 | 1,043 | 1,944 | 31.8 | 11.7 | 344.2 | 293.0 | 60.4% | 68.2% |
| 60% | 11,505 | 993 | 1,128 | 2,139 | 28.6 | 10.2 | 312.7 | 309.3 | 64.4% | 72.0% |
| 70% | 12,782 | 1,098 | 1,197 | 2,311 | 26.2 | 9.3 | 290.3 | 321.2 | 67.4% | 74.7% |
| 80% | 13,965 | 1,187 | 1,254 | 2,457 | 24.2 | 8.5 | 271.1 | 330.9 | 69.9% | 76.9% |
| 90% | 15,041 | 1,272 | 1,303 | 2,589 | 22.6 | 7.8 | 253.5 | 338.5 | 71.9% | 78.6% |
| 100% | 16,023 | 1,350 | 1,344 | 2,707 | 21.3 | 7.4 | 239.1 | 343.6 | 73.4% | 79.8% |

COL, colonoscopy; CRC, colorectal cancer; FIT, fecal immunochemical test; HSgFOBT, high-sensitivity guaiac-based fecal occult blood test; LY, life-years; LYG, life-years gained; mt-sDNA, multitarget stool DNA test.

**Table S3. Outcomes and efficiency ratios of LYG relative to number of colonoscopies assuming perfect (100%) adherence.** Results are ordered by total colonoscopies. Results shown are per 1000 individuals free of diagnosed colorectal cancer at age 40 and screened starting at age 45 or 50 and ending at age 75 or 80 receiving biennial or triennial mt-sDNA, annual or biennial FIT, and annual or biennial HSgFOBT. Bold row is the model-recommended strategy.

| Screening Strategy | Stool Tests | Total COL | LYG | Complications | CRC Deaths Averted | ΔCOL | ΔLYG | Efficiency Ratio (ΔCOL/ΔLYG) | Detail |
|---|---|---|---|---|---|---|---|---|---|
| FIT 50-75, 2 | 9,447 | 1,480 | 273.5 | 9 | 23.7 | ND | ND | ND | Efficient |
| FIT 50-80, 2 | 10,790 | 1,598 | 289.3 | 11 | 26.0 | 118.0 | 15.7 | 7.5 | Efficient |
| FIT 45-75, 2 | 11,670 | 1,686 | 300.2 | 10 | 25.3 | 87.8 | 10.9 | 8.0 | Near Efficient |
| HSgFOBT 50-75, 2 | 8,627 | 1,758 | 278.5 | 10 | 24.3 | ND | ND | ND | Strongly Dominated |
| FIT 45-80, 2 | 12,562 | 1,763 | 310.9 | 11 | 26.8 | 165.1 | 21.6 | 7.6 | Efficient |
| HSgFOBT 50-80, 2 | 9,831 | 1,905 | 293.5 | 11 | 26.5 | ND | ND | ND | Strongly Dominated |
| mt-sDNA 50-75, 3 | 6,076 | 1,957 | 300.0 | 11 | 26.4 | ND | ND | ND | Strongly Dominated |
| HSgFOBT 45-75, 2 | 10,608 | 2,025 | 305.8 | 10 | 25.9 | ND | ND | ND | Strongly Dominated |
| FIT 50-75, 1 | 15,856 | 2,037 | 318.1 | 11 | 27.9 | ND | ND | ND | Weakly Dominated |
| mt-sDNA 50-80, 3 | 6,676 | 2,068 | 309.5 | 12 | 27.9 | ND | ND | ND | Strongly Dominated |
| HSgFOBT 45-80, 2 | 11,407 | 2,120 | 315.4 | 11 | 27.4 | ND | ND | ND | Strongly Dominated |
| FIT 50-80, 1 | 17,642 | 2,159 | 328.2 | 13 | 29.5 | ND | ND | ND | Weakly Dominated |
| mt-sDNA 45-75, 3 | 7,213 | 2,196 | 320.9 | 11 | 27.3 | ND | ND | ND | Strongly Dominated |
| mt-sDNA 50-75, 2 | 7,634 | 2,277 | 319.3 | 12 | 28.1 | ND | ND | ND | Strongly Dominated |
| FIT 45-75, 1 | 19,182 | 2,285 | 340.7 | 12 | 28.9 | 522.1 | 29.8 | 17.5 | Near Efficient |
| mt-sDNA 45-80, 3 | 7,856 | 2,313 | 331.5 | 12 | 28.9 | ND | ND | ND | Strongly Dominated |

| | | | | | | | | | |
|---|---|---|---|---|---|---|---|---|---|
| HSgFOBT 50-75, 1 | 13,337 | 2,388 | 320.6 | 12 | 28.2 | ND | ND | ND | Strongly Dominated |
| **FIT 45-80, 1** | **20,958** | **2,405** | **351.1** | **13** | **30.5** | **642.3** | **40.2** | **16.0** | **Efficient** |
| mt-sDNA 50-80, 2 | 8,644 | 2,442 | 330.1 | 13 | 29.9 | ND | ND | ND | Strongly Dominated |
| HSgFOBT 50-80, 1 | 14,792 | 2,544 | 330.0 | 13 | 29.7 | ND | ND | ND | Strongly Dominated |
| mt-sDNA 45-75, 2 | 9,382 | 2,599 | 344.3 | 12 | 29.4 | ND | ND | ND | Strongly Dominated |
| HSgFOBT 45-75, 1 | 16,023 | 2,707 | 343.6 | 12 | 29.2 | ND | ND | ND | Strongly Dominated |
| mt-sDNA 45-80, 2 | 10,046 | 2,712 | 352.8 | 13 | 30.7 | 306.6 | 1.7 | 180.3 | Efficient |
| HSgFOBT 45-80, 1 | 17,489 | 2,859 | 352.9 | 13 | 30.7 | 146.9 | 0.1 | 1093.6 | Efficient |

COL, colonoscopy; CRC, colorectal cancer; FIT, fecal immunochemical test; HSgFOBT, high-sensitivity guaiac-based fecal occult blood test; LYG, life-years gained; mt-sDNA, multitarget stool DNA test; ND, indicates an efficiency ratio is not defined because the strategy is not efficient or near-efficient.

**Table S4. Outcomes and efficiency ratios of LYG relative to number of colonoscopies at base-case imperfect adherence rates of 40% FIT vs 34% HSgFOBT vs 70% mt-sDNA.** Results are ordered by total colonoscopies. Results shown are per 1000 individuals free of diagnosed colorectal cancer at age 40 and screened starting at age 45 or 50 and ending at age 75 or 80 receiving biennial or triennial mt-sDNA, annual or biennial FIT, and annual or biennial HSgFOBT. Gray highlight indicates shift from dominated to efficient or near-efficient from 100% adherence assumption. Italics indicates shift from efficient or near-efficient to dominated from 100% adherence assumption. Bold row is the model-recommended strategy.

| Screening Strategy | Stool Tests | Total COL | LYG | Complications | CRC Deaths Averted | ΔCOL | ΔLYG | Efficiency Ratio (ΔCOL/ΔLYG) | Detail |
|---|---|---|---|---|---|---|---|---|---|
| FIT 50-75, 2 | 5,856 | 1,056 | 210.9 | 8 | 18.5 | ND | ND | ND | Efficient |
| FIT 50-80, 2 | 6,581 | 1,132 | 225.0 | 8 | 20.5 | 76.7 | 14.2 | 5.4 | Efficient |
| HSgFOBT 50-75, 2 | 5,052 | 1,160 | 203.6 | 8 | 18.0 | ND | ND | ND | Strongly Dominated |
| FIT 45-75, 2 | 7,073 | 1,190 | 232.9 | 8 | 19.7 | 57.4 | 7.9 | 7.3 | Near Efficient |
| HSgFOBT 50-80, 2 | 5,674 | 1,250 | 215.8 | 9 | 19.8 | ND | ND | ND | Strongly Dominated |
| FIT 45-80, 2 | 7,798 | 1,264 | 245.6 | 9 | 21.5 | 131.1 | 20.6 | 6.4 | Efficient |
| FIT 50-75, 1 | 7,672 | 1,269 | 239.4 | 8 | 21.1 | ND | ND | ND | Strongly Dominated |
| HSgFOBT 45-75, 2 | 6,089 | 1,325 | 224.3 | 8 | 19.1 | ND | ND | ND | Strongly Dominated |
| FIT 50-80, 1 | 8,622 | 1,358 | 253.2 | 10 | 23.1 | 94.6 | 7.6 | 12.5 | Near Efficient |
| HSgFOBT 50-75, 1 | 6,281 | 1,371 | 227.4 | 8 | 20.3 | ND | ND | ND | Strongly Dominated |
| HSgFOBT 45-80, 2 | 6,709 | 1,413 | 238.0 | 9 | 21.0 | ND | ND | ND | Strongly Dominated |
| FIT 45-75, 1 | 9,293 | 1,433 | 262.7 | 9 | 22.3 | 169.2 | 17.1 | 9.9 | Near Efficient |
| HSgFOBT 50-80, 1 | 7,042 | 1,473 | 239.8 | 10 | 22.1 | ND | ND | ND | Strongly Dominated |
| FIT 45-80, 1 | 10,223 | 1,519 | 275.3 | 10 | 24.2 | 255.2 | 29.7 | 8.6 | Efficient |
| HSgFOBT 45-75, 1 | 7,573 | 1,564 | 251.7 | 9 | 21.6 | ND | ND | ND | Strongly Dominated |
| *HSgFOBT 45-80, 1* | *8,330* | *1,666* | *265.5* | *10* | *23.4* | *ND* | *ND* | *ND* | *Strongly Dominated* |
| mt-sDNA 50-75, 3 | 5,397 | 1,813 | 285.1 | 10 | 25.2 | ND | ND | ND | Weakly Dominated |

| | | | | | | | | | |
|---|---|---|---|---|---|---|---|---|---|
| mt-sDNA 50-80, 3 | 6,010 | 1,929 | 296.9 | 11 | 26.9 | ND | ND | ND | Weakly Dominated |
| mt-sDNA 45-75, 3 | 6,485 | 2,043 | 309.0 | 10 | 26.3 | 524.5 | 33.7 | 15.5 | Near Efficient |
| mt-sDNA 50-75, 2 | 6,725 | 2,096 | 307.2 | 11 | 27.1 | ND | ND | ND | Strongly Dominated |
| mt-sDNA 45-80, 3 | 7,090 | 2,156 | 319.7 | 12 | 28.0 | 636.7 | 44.4 | 14.3 | Efficient |
| mt-sDNA 50-80, 2 | 7,479 | 2,224 | 317.3 | 13 | 28.7 | ND | ND | ND | Strongly Dominated |
| mt-sDNA 45-75, 2 | 8,092 | 2,362 | 331.2 | 11 | 28.3 | 206.4 | 11.5 | 18.0 | Near Efficient |
| **mt-sDNA 45-80, 2** | **8,826** | **2,495** | **339.8** | **13** | **29.7** | **339.9** | **20.1** | **17.0** | **Efficient** |

COL, colonoscopy; CRC, colorectal cancer; FIT, fecal immunochemical test; HSgFOBT, high-sensitivity guaiac-based fecal occult blood testLYG, life-years gained; mt-sDNA, multitarget stool DNA test; ND, indicates an efficiency ratio is not defined because the strategy is not efficient or near-efficient.

**Table S5. Incremental efficiency ratios for number of colonoscopies at a fixed screening window of 50–75 or 45–75 assuming perfect (100%) adherence, base-case imperfect adherence rates of 40% FIT vs 34% HSgFOBT vs 70% mt-sDNA, adherence of 50% FIT vs 43% HSgFOBT vs 70% mt-sDNA, or adherence of 60% FIT vs 52% HSgFOBT vs 70% mt-sDNA.** Results are ordered by total colonoscopies. Results shown are per 1000 individuals free of diagnosed colorectal cancer receiving biennial or triennial mt-sDNA, annual or biennial FIT, and annual or biennial HSgFOBT.

| Screen Window | Adherence Scenario | Strategies | Total COLs | LYG | ΔCOL | ΔLYG | Efficiency Ratio (ΔCOL/ΔLYG) | Detail |
|---|---|---|---|---|---|---|---|---|
| 50–75 | 100% FIT/HSgFOBT/mt-sDNA | FIT 50-75, 2 | 1,480 | 273.5 | -- | -- | -- | Efficient |
| | | HSgFOBT 50-75, 2 | 1,758 | 278.5 | ND | ND | ND | Weakly Dominated |
| | | mt-sDNA 50-75, 3 | 1,957 | 300.0 | ND | ND | ND | Weakly Dominated |
| | | FIT 50-75, 1 | 2,037 | 318.1 | 557.0 | 44.6 | 12.5 | Efficient |
| | | mt-sDNA 50-75, 2 | 2,277 | 319.3 | 239.9 | 1.2 | 196.6 | Near Efficient |
| | | HSgFOBT 50-75, 1 | 2,388 | 320.6 | 351.2 | 2.5 | 142.8 | Efficient |
| 50–75 | 40% FIT/34% HSgFOBT/70% mt-sDNA | FIT 50-75, 2 | 1,056 | 210.9 | -- | -- | -- | Efficient |
| | | HSgFOBT 50-75, 2 | 1,160 | 203.6 | ND | ND | ND | Strongly Dominated |
| | | FIT 50-75, 1 | 1,269 | 239.4 | 213.4 | 28.5 | 7.5 | Efficient |
| | | HSgFOBT 50-75, 1 | 1,371 | 227.4 | ND | ND | ND | Strongly Dominated |
| | | mt-sDNA 50-75, 3 | 1,813 | 285.1 | 543.8 | 45.7 | 11.9 | Efficient |
| | | mt-sDNA 50-75, 2 | 2,096 | 307.2 | 282.8 | 22.1 | 12.8 | Efficient |
| 50–75 | 50% FIT/43% HSgFOBT/70% mt-sDNA | FIT 50-75, 2 | 1,164 | 229.3 | -- | -- | -- | Efficient |
| | | HSgFOBT 50-75, 2 | 1,298 | 223.3 | ND | ND | ND | Strongly Dominated |
| | | FIT 50-75, 1 | 1,444 | 263.0 | 280.0 | 33.8 | 8.3 | Efficient |
| | | HSgFOBT 50-75, 1 | 1,574 | 252.2 | ND | ND | ND | Strongly Dominated |
| | | mt-sDNA 50-75, 3 | 1,813 | 285.1 | 368.6 | 22.1 | 16.7 | Near Efficient |
| | | mt-sDNA 50-75, 2 | 2,096 | 307.2 | 651.4 | 44.2 | 14.7 | Efficient |
| 50–75 | | FIT 50-75, 2 | 1,256 | 243.0 | -- | -- | -- | Efficient |

| | | | | | | | | |
|---|---|---|---|---|---|---|---|---|
| | 60% FIT/52% HSgFOBT/70% mt-sDNA | HSgFOBT 50-75, 2 | 1,408 | 237.3 | ND | ND | ND | Strongly Dominated |
| | | FIT 50-75, 1 | 1,594 | 279.5 | 338.5 | 36.5 | 9.3 | Efficient |
| | | HSgFOBT 50-75, 1 | 1,747 | 271.4 | ND | ND | ND | Strongly Dominated |
| | | mt-sDNA 50-75, 3 | 1,813 | 285.1 | ND | ND | ND | Weakly Dominated |
| | | mt-sDNA 50-75, 2 | 2,096 | 307.2 | 501.7 | 27.7 | 18.1 | Efficient |
| 45–75 | 100% FIT/HSgFOBT/mt-sDNA | FIT 45-75, 2 | 1,686 | 300.2 | -- | -- | -- | Efficient |
| | | HSgFOBT 45-75, 2 | 2,025 | 305.8 | ND | ND | ND | Weakly Dominated |
| | | mt-sDNA 45-75, 3 | 2,196 | 320.9 | ND | ND | ND | Weakly Dominated |
| | | FIT 45-75, 1 | 2,285 | 340.7 | 599 | 40 | 14.8 | Efficient |
| | | mt-sDNA 45-75, 2 | 2,599 | 344.3 | 314 | 4 | 86.0 | Efficient |
| | | HSgFOBT 45-75, 1 | 2,707 | 343.6 | ND | ND | ND | Strongly Dominated |
| 45–75 | 40% FIT/34% HSgFOBT/70% mt-sDNA | FIT 45-75, 2 | 1,190 | 232.9 | -- | -- | -- | Efficient |
| | | HSgFOBT 45-75, 2 | 1,325 | 224.3 | ND | ND | ND | Strongly Dominated |
| | | FIT 45-75, 1 | 1,433 | 262.7 | 242.9 | 29.8 | 8.1 | Efficient |
| | | HSgFOBT 45-75, 1 | 1,564 | 251.7 | ND | ND | ND | Strongly Dominated |
| | | mt-sDNA 45-75, 3 | 2,043 | 309.0 | 610.5 | 46.3 | 13.2 | Efficient |
| | | mt-sDNA 45-75, 2 | 2,362 | 331.2 | 318.6 | 22.2 | 14.4 | Efficient |
| 45–75 | 50% FIT/43% HSgFOBT/70% mt-sDNA | FIT 45-75, 2 | 1,314 | 251.6 | -- | -- | -- | Efficient |
| | | HSgFOBT 45-75, 2 | 1,479 | 245.7 | ND | ND | ND | Strongly Dominated |
| | | FIT 45-75, 1 | 1,628 | 286.5 | 313.8 | 34.9 | 9.0 | Efficient |
| | | HSgFOBT 45-75, 1 | 1,792 | 278.1 | ND | ND | ND | Strongly Dominated |
| | | mt-sDNA 45-75, 3 | 2,043 | 309.0 | 415.5 | 22.6 | 18.4 | Near Efficient |
| | | mt-sDNA 45-75, 2 | 2,362 | 331.2 | 734.1 | 44.7 | 16.4 | Efficient |
| 45–75 | 60% FIT/52% HSgFOBT/70% mt-sDNA | FIT 45-75, 2 | 1,413 | 266.7 | -- | -- | -- | Efficient |
| | | HSgFOBT 45-75, 2 | 1,603 | 260.8 | ND | ND | ND | Strongly Dominated |
| | | FIT 45-75, 1 | 1,795 | 304.3 | 382.0 | 37.6 | 10.2 | Efficient |

| | | | | | | | | |
|---|---|---|---|---|---|---|---|---|
| | | HSgFOBT 45-75, 1 | 1,987 | 295.8 | ND | ND | ND | Strongly Dominated |
| | | mt-sDNA 45-75, 3 | 2,043 | 309.0 | ND | ND | ND | Weakly Dominated |
| | | mt-sDNA 45-75, 2 | 2,362 | 331.2 | 567.0 | 27.0 | 21.0 | Efficient |

COL, colonoscopy; CRC, colorectal cancer; FIT, fecal immunochemical test; HSgFOBT, high-sensitivity guaiac-based fecal occult blood test; LYG, life-years gained; mt-sDNA, multitarget stool DNA test; ND, indicates an efficiency ratio is not defined because the strategy is not efficient or near-efficient.

**Table S6. Outcomes and efficiency ratios based on patient hours related to the screening process assuming perfect (100%) adherence.** Results are ordered by patient hours. Results shown are per 1000 individuals free of diagnosed colorectal cancer at age 40 and screened starting at age 45 or 50 and ending at age 75 or 80 receiving biennial or triennial mt-sDNA, annual or biennial FIT, and annual or biennial HSgFOBT. Bold row is the model-recommended strategy.

| Screening Strategy | Stool Tests | Patient Hours | LYG | Complications | CRC Deaths Averted | ΔHours | ΔLYG | Efficiency Ratio (ΔHours/ΔLYG) | Detail |
|---|---|---|---|---|---|---|---|---|---|
| FIT 50-75, 2 | 9,447 | 34,175 | 273.5 | 9 | 23.7 | ND | ND | ND | Efficient |
| FIT 50-80, 2 | 10,790 | 37,549 | 289.3 | 11 | 26.0 | 3,374.5 | 15.7 | 214.3 | Near Efficient |
| mt-sDNA 50-75, 3 | 6,076 | 38,613 | 300.0 | 11 | 26.4 | 4,438.6 | 26.4 | 167.9 | Efficient |
| FIT 45-75, 2 | 11,670 | 39,751 | 300.2 | 10 | 25.3 | 1,137.6 | 0.2 | 4,795.7 | Near Efficient |
| mt-sDNA 50-80, 3 | 6,676 | 41,102 | 309.5 | 12 | 27.9 | 2,488.6 | 9.5 | 262.4 | Near Efficient |
| FIT 45-80, 2 | 12,562 | 41,967 | 310.9 | 11 | 26.8 | 3,353.4 | 10.9 | 307.0 | Near Efficient |
| mt-sDNA 45-75, 3 | 7,213 | 43,579 | 320.9 | 11 | 27.3 | 4,966.1 | 20.9 | 237.7 | Efficient |
| mt-sDNA 50-75, 2 | 7,634 | 45,388 | 319.3 | 12 | 28.1 | ND | ND | ND | Strongly Dominated |
| mt-sDNA 45-80, 3 | 7,856 | 46,223 | 331.5 | 12 | 28.9 | 2,643.3 | 10.6 | 249.8 | Efficient |
| HSgFOBT 50-75, 2 | 8,627 | 46,489 | 278.5 | 10 | 24.3 | ND | ND | ND | Strongly Dominated |
| mt-sDNA 50-80, 2 | 8,644 | 49,204 | 330.1 | 13 | 29.9 | ND | ND | ND | Strongly Dominated |
| FIT 50-75, 1 | 15,856 | 49,719 | 318.1 | 11 | 27.9 | ND | ND | ND | Strongly Dominated |
| HSgFOBT 50-80, 2 | 9,831 | 51,405 | 293.5 | 11 | 26.5 | ND | ND | ND | Strongly Dominated |
| mt-sDNA 45-75, 2 | 9,382 | 52,318 | 344.3 | 12 | 29.4 | 6,095.5 | 12.9 | 473.1 | Near Efficient |
| FIT 50-80, 1 | 17,642 | 53,588 | 328.2 | 13 | 29.5 | ND | ND | ND | Strongly Dominated |
| HSgFOBT 45-75, 2 | 10,608 | 54,756 | 305.8 | 10 | 25.9 | ND | ND | ND | Strongly Dominated |

| | | | | | | | | | |
|---|---|---|---|---|---|---|---|---|---|
| **mt-sDNA 45-80, 2** | **10,046** | **54,906** | **352.8** | **13** | **30.7** | **8,683.5** | **21.4** | **406.6** | **Efficient** |
| FIT 45-75, 1 | 19,182 | 57,047 | 340.7 | 12 | 28.9 | ND | ND | ND | Strongly Dominated |
| HSgFOBT 45-80, 2 | 11,407 | 57,999 | 315.4 | 11 | 27.4 | ND | ND | ND | Strongly Dominated |
| FIT 45-80, 1 | 20,958 | 60,879 | 351.1 | 13 | 30.5 | ND | ND | ND | Strongly Dominated |
| HSgFOBT 50-75, 1 | 13,337 | 66,216 | 320.6 | 12 | 28.2 | ND | ND | ND | Strongly Dominated |
| HSgFOBT 50-80, 1 | 14,792 | 71,754 | 330.0 | 13 | 29.7 | ND | ND | ND | Strongly Dominated |
| HSgFOBT 45-75, 1 | 16,023 | 76,709 | 343.6 | 12 | 29.2 | ND | ND | ND | Strongly Dominated |
| HSgFOBT 45-80, 1 | 17,489 | 82,207 | 352.9 | 13 | 30.7 | 27,301.0 | 0.1 | 203,188.5 | Efficient |

COL, colonoscopy; CRC, colorectal cancer; FIT, fecal immunochemical test; HSgFOBT, high-sensitivity guaiac-based fecal occult blood test; LYG, life-years gained; mt-sDNA, multitarget stool DNA test; ND, indicates an efficiency ratio is not defined because the strategy is not efficient or near-efficient.

**Table S7. Outcomes and efficiency ratios based on patient hours related to the screening process at base-case imperfect adherence rates of 40% FIT vs 34% HSgFOBT vs 70% mt-sDNA.** Results are ordered by patient hours. Results shown are per 1000 individuals free of diagnosed colorectal cancer at age 40 and screened starting at age 45 or 50 and ending at age 75 or 80 receiving biennial or triennial mt-sDNA, annual or biennial FIT, and annual or biennial HSgFOBT. Italics indicates shift from efficient or near-efficient to dominated from 100% adherence assumption. Bold row is the model-recommended strategy.

| Screening Strategy | Stool Tests | Patient Hours | LYG | Complications | CRC Deaths Averted | ΔHours | ΔLYG | Efficiency Ratio (ΔHours/ΔLYG) | Detail |
|---|---|---|---|---|---|---|---|---|---|
| FIT 50-75, 2 | 5,856 | 23,604 | 210.9 | 8 | 18.5 | ND | ND | ND | Efficient |
| FIT 50-80, 2 | 6,581 | 25,652 | 225.0 | 8 | 20.5 | 2,047.7 | 14.2 | 144.4 | Efficient |
| FIT 45-75, 2 | 7,073 | 26,968 | 232.9 | 8 | 19.7 | 1315.5 | 7.9 | 166.9 | Near Efficient |
| FIT 50-75, 1 | 7,672 | 28,930 | 239.4 | 8 | 21.1 | ND | ND | ND | Weakly Dominated |
| FIT 45-80, 2 | 7,798 | 28,988 | 245.6 | 9 | 21.5 | 3,335.4 | 20.6 | 162.0 | Efficient |
| HSgFOBT 50-75, 2 | 5,052 | 29,517 | 203.6 | 8 | 18.0 | ND | ND | ND | Strongly Dominated |
| FIT 50-80, 1 | 8,622 | 31,426 | 253.2 | 10 | 23.1 | ND | ND | ND | Weakly Dominated |
| HSgFOBT 50-80, 2 | 5,674 | 32,313 | 215.8 | 9 | 19.8 | ND | ND | ND | Strongly Dominated |
| FIT 45-75, 1 | 9,293 | 33,189 | 262.7 | 9 | 22.3 | ND | ND | ND | Weakly Dominated |
| HSgFOBT 45-75, 2 | 6,089 | 34,269 | 224.3 | 8 | 19.1 | ND | ND | ND | Strongly Dominated |
| HSgFOBT 50-75, 1 | 6,281 | 35,449 | 227.4 | 8 | 20.3 | ND | ND | ND | Strongly Dominated |
| mt-sDNA 50-75, 3 | 5,397 | 35,555 | 285.1 | 10 | 25.2 | 6,567.4 | 39.4 | 166.6 | Efficient |
| FIT 45-80, 1 | 10,223 | 35,625 | 275.3 | 10 | 24.2 | ND | ND | ND | Strongly Dominated |

| | | | | | | | | | |
|---|---|---|---|---|---|---|---|---|---|
| HSgFOBT 45-80, 2 | 6,709 | 37,019 | 238.0 | 9 | 21.0 | ND | ND | ND | Strongly Dominated |
| mt-sDNA 50-80, 3 | 6,010 | 38,145 | 296.9 | 11 | 26.9 | 2,590.4 | 11.8 | 218.9 | Near Efficient |
| HSgFOBT 50-80, 1 | 7,042 | 38,730 | 239.8 | 10 | 22.1 | ND | ND | ND | Strongly Dominated |
| mt-sDNA 45-75, 3 | 6,485 | 40,350 | 309.0 | 10 | 26.3 | 4,794.9 | 24.0 | 199.9 | Efficient |
| HSgFOBT 45-75, 1 | 7,573 | 41,161 | 251.7 | 9 | 21.6 | ND | ND | ND | Strongly Dominated |
| mt-sDNA 50-75, 2 | 6,725 | 41,531 | 307.2 | 11 | 27.1 | ND | ND | ND | Strongly Dominated |
| mt-sDNA 45-80, 3 | 7,090 | 42,874 | 319.7 | 12 | 28.0 | 2,524.5 | 10.7 | 236.4 | Efficient |
| *HSgFOBT 45-80, 1* | *8,330* | *44,417* | *265.5* | *10* | *23.4* | *ND* | *ND* | *ND* | *Strongly Dominated* |
| mt-sDNA 50-80, 2 | 7,479 | 44,481 | 317.3 | 13 | 28.7 | ND | ND | ND | Strongly Dominated |
| mt-sDNA 45-75, 2 | 8,092 | 47,162 | 331.2 | 11 | 28.3 | 4,288.0 | 11.5 | 373.0 | Near Efficient |
| **mt-sDNA 45-80, 2** | **8,826** | **50,170** | **339.8** | **13** | **29.7** | **7,296.0** | **20.1** | **363.9** | **Efficient** |

COL, colonoscopy; CRC, colorectal cancer; FIT, fecal immunochemical test; HSgFOBT, high-sensitivity guaiac-based fecal occult blood test; LYG, life-years gained; mt-sDNA, multitarget stool DNA test; ND, indicates an efficiency ratio is not defined because the strategy is not efficient or near-efficient.

**Table S8. Incremental efficiency ratios for patient hours related to the screening process at a fixed screening window of 50–75 or 45–75 assuming perfect (100%) adherence, base-case imperfect adherence rates of 40% FIT vs 34% HSgFOBT vs 70% mt-sDNA, 50% FIT vs 43% HSgFOBT vs 70% mt-sDNA adherence, or 60% FIT vs 52% HSgFOBT vs 70% mt-sDNA adherence.** Results are ordered by patient hours. Results shown are per 1000 individuals free of diagnosed colorectal cancer receiving biennial or triennial mt-sDNA, annual or biennial FIT, and annual or biennial HSgFOBT.

| Screen Window | Adherence Scenario | Strategies | Patient Hours | LYG | ΔHours | ΔLYG | Efficiency Ratio (ΔHours/ΔLYG) | Detail |
|---|---|---|---|---|---|---|---|---|
| 50–75 | 100% FIT/HSgFOBT/mt-sDNA | FIT 50-75, 2 | 34,175 | 273.5 | -- | -- | -- | Efficient |
| | | mt-sDNA 50-75, 3 | 38,613 | 300.0 | 4,438.6 | 26.4 | 167.9 | Efficient |
| | | mt-sDNA 50-75, 2 | 45,388 | 319.3 | 6,774.4 | 19.4 | 350.0 | Efficient |
| | | HSgFOBT 50-75, 2 | 46,489 | 278.5 | ND | ND | ND | Strongly Dominated |
| | | FIT 50-75, 1 | 49,719 | 318.1 | ND | ND | ND | Strongly Dominated |
| | | HSgFOBT 50-75, 1 | 66,216 | 320.6 | 20,828.6 | 1.2 | 16,807.5 | Efficient |
| 50–75 | 40% FIT/34% HSgFOBT/70% mt-sDNA | FIT 50-75, 2 | 23,604 | 210.9 | -- | -- | -- | Efficient |
| | | FIT 50-75, 1 | 28,930 | 239.4 | 5,325.2 | 28.5 | 186.6 | Near Efficient |
| | | HSgFOBT 50-75, 2 | 29,517 | 203.6 | ND | ND | ND | Strongly Dominated |
| | | HSgFOBT 50-75, 1 | 35,449 | 227.4 | ND | ND | ND | Strongly Dominated |
| | | mt-sDNA 50-75, 3 | 35,555 | 285.1 | 11,950.6 | 74.2 | 161.1 | Efficient |
| | | mt-sDNA 50-75, 2 | 41,531 | 307.2 | 5,976.0 | 22.1 | 269.8 | Efficient |
| 50–75 | 50% FIT/43% HSgFOBT/70% mt-sDNA | FIT 50-75, 2 | 26,277 | 229.3 | -- | -- | -- | Efficient |
| | | HSgFOBT 50-75, 2 | 33,287 | 223.3 | ND | ND | ND | Strongly Dominated |
| | | FIT 50-75, 1 | 33,394 | 263.0 | ND | ND | ND | Weakly Dominated |
| | | mt-sDNA 50-75, 3 | 35,555 | 285.1 | 9,278.0 | 55.8 | 166.2 | Efficient |
| | | HSgFOBT 50-75, 1 | 41,203 | 252.2 | ND | ND | ND | Strongly Dominated |
| | | mt-sDNA 50-75, 2 | 41,531 | 307.2 | 5,976.0 | 22.1 | 269.8 | Efficient |
| 50–75 | | FIT 50-75, 2 | 28,514 | 243.0 | -- | -- | -- | Efficient |

| | | | | | | | | |
|---|---|---|---|---|---|---|---|---|
| | 60% FIT/52% HSgFOBT/70% mt-sDNA | mt-sDNA 50-75, 3 | 35,555 | 285.1 | 7,041.0 | 42.1 | 167.3 | Efficient |
| | | HSgFOBT 50-75, 2 | 36,402 | 237.3 | ND | ND | ND | Strongly Dominated |
| | | FIT 50-75, 1 | 37,331 | 279.5 | ND | ND | ND | Strongly Dominated |
| | | mt-sDNA 50-75, 2 | 41,531 | 307.2 | 5,976.0 | 22.1 | 269.8 | Efficient |
| | | HSgFOBT 50-75, 1 | 46,304 | 271.4 | ND | ND | ND | Strongly Dominated |
| 45–75 | 100% FIT/HSgFOBT/mt-sDNA | FIT 45-75, 2 | 39,751 | 300.2 | -- | -- | -- | Efficient |
| | | mt-sDNA 45-75, 3 | 43,579 | 320.9 | 3,828.5 | 20.7 | 185.3 | Efficient |
| | | mt-sDNA 45-75, 2 | 52,318 | 344.3 | 8,738.9 | 23.5 | 372.4 | Efficient |
| | | HSgFOBT 45-75, 2 | 54,756 | 305.8 | ND | ND | ND | Strongly Dominated |
| | | FIT 45-75, 1 | 57,047 | 340.7 | ND | ND | ND | Strongly Dominated |
| | | HSgFOBT 45-75, 1 | 76,709 | 343.6 | ND | ND | ND | Strongly Dominated |
| 45–75 | 40% FIT/34% HSgFOBT/70% mt-sDNA | FIT 45-75, 2 | 26,968 | 232.9 | -- | -- | -- | Efficient |
| | | FIT 45-75, 1 | 33,189 | 262.7 | ND | ND | ND | Weakly Dominated |
| | | HSgFOBT 45-75, 2 | 34,269 | 224.3 | ND | ND | ND | Strongly Dominated |
| | | mt-sDNA 45-75, 3 | 40,350 | 309.0 | 13,382.2 | 76.1 | 175.8 | Efficient |
| | | HSgFOBT 45-75, 1 | 41,161 | 251.7 | ND | ND | ND | Strongly Dominated |
| | | mt-sDNA 45-75, 2 | 47,162 | 331.2 | 6,812.5 | 22.2 | 307.3 | Efficient |
| 45–75 | 50% FIT/43% HSgFOBT/70% mt-sDNA | FIT 45-75, 2 | 30,064 | 251.6 | -- | -- | -- | Efficient |
| | | FIT 45-75, 1 | 38,289 | 286.5 | ND | ND | ND | Weakly Dominated |
| | | HSgFOBT 45-75, 2 | 38,586 | 245.7 | ND | ND | ND | Strongly Dominated |
| | | mt-sDNA 45-75, 3 | 40,350 | 309.0 | 10,286.4 | 57.5 | 179.0 | Efficient |
| | | mt-sDNA 45-75, 2 | 47,162 | 331.2 | 6,812.5 | 22.2 | 307.3 | Efficient |
| | | HSgFOBT 45-75, 1 | 47,792 | 278.1 | ND | ND | ND | Strongly Dominated |
| 45–75 | | FIT 45-75, 2 | 32,586 | 266.7 | -- | -- | -- | Efficient |
| | | mt-sDNA 45-75, 3 | 40,350 | 309.0 | 7,763.5 | 42.4 | 183.4 | Efficient |

| | | | | | | | | |
|---|---|---|---|---|---|---|---|---|
| | 60% FIT/52% HSgFOBT/70% mt-sDNA | HSgFOBT 45-75, 2 | 42,172 | 260.8 | ND | ND | ND | Strongly Dominated |
| | | FIT 45-75, 1 | 42,817 | 304.3 | ND | ND | ND | Strongly Dominated |
| | | mt-sDNA 45-75, 2 | 47,162 | 331.2 | 6,812.5 | 22.2 | 307.3 | Efficient |
| | | HSgFOBT 45-75, 1 | 53,694 | 295.8 | ND | ND | ND | Strongly Dominated |

COL, colonoscopy; CRC, colorectal cancer; FIT, fecal immunochemical test; HSgFOBT, high-sensitivity guaiac-based fecal occult blood test; LYG, life-years gained; mt-sDNA, multitarget stool DNA test; ND, indicates an efficiency ratio is not defined because the strategy is not efficient or near-efficient.

**Table S9. Predicted outcomes per 1000 individuals screened from ages 45–75 compared with no screening**. Individuals were randomly assigned numbers of mt-sDNA (max=11 triennial tests during the screening window) or FIT (max=31 annual tests during the screening window).

| Screening strategy | Randomly Assigned Number of Tests | Total Stool Tests | Total COLs | CRC Cases | CRC Deaths | LY with CRC | LYG | Incremental COL/ Incremental LYG vs FIT | Incidence Reduction | Mortality Reduction |
|---|---|---|---|---|---|---|---|---|---|---|
| mt-sDNA, 45-75 | Up to 1 | 831 | 421 | 67.8 | 29.5 | 600.1 | 69.6 | 6.5 | 15.6% | 19.4% |
| FIT, 45-75 | Up to 1 | 825 | 245 | 73.8 | 32.5 | 631.1 | 42.4 | | 8.1% | 11.2% |
| mt-sDNA, 45-75 | Up to 5 | 3,555 | 1,336 | 41.0 | 15.8 | 442.6 | 222.9 | 7.9 | 48.9% | 56.8% |
| FIT, 45-75 | Up to 5 | 3,790 | 746 | 55.6 | 22.3 | 558.8 | 148.0 | | 30.7% | 39.2% |
| mt-sDNA, 45-75 | Up to 11 | 7,119 | 2,182 | 26.0 | 9.2 | 289.3 | 320.7 | 10.2 | 67.7% | 74.9% |
| FIT, 45-75 | Up to 11 | 7,658 | 1,254 | 41.0 | 15.2 | 464.0 | 229.8 | | 49.0% | 58.5% |

COL, colonoscopy; CRC, colorectal cancer; FIT, fecal immunochemical test; LY, life-years; LYG, life-years gained; mt-sDNA, multitarget stool DNA test.

**Table S10. Deep-C (clinicaltrials.gov identifier, NCT01397747) sensitivity analysis for screening outcomes per 1000 individuals by adherence rate for triennial mt-sDNA and annual FIT in individuals free of diagnosed colorectal cancer at age 40 and screened between ages 50–75 years or 45–75 years.**

| Screening Strategy and Adherence Rate | Stool Tests | Follow-up COLs | Surveillance COLs | Total COLs | CRC Cases | CRC Deaths | LY with CRC | LYG | Incidence Reduction | Mortality Reduction |
|---|---|---|---|---|---|---|---|---|---|---|
| No screening | 0 | 0 | 0 | 80 | 80.3 | 36.6 | 646.0 | 0.0 | 0.0% | 0.0% |
| mt-sDNA 50-75, 3 | | | | | | | | | | |
| 10% | 1,801 | 248 | 468 | 765 | 56.1 | 23.8 | 515.5 | 140.9 | 30.2% | 35.0% |
| 20% | 2,968 | 387 | 700 | 1,123 | 45.2 | 18.3 | 448.2 | 204.0 | 43.7% | 50.1% |
| 30% | 3,795 | 478 | 834 | 1,341 | 39.3 | 15.4 | 406.3 | 237.9 | 51.1% | 58.0% |
| 40% | 4,415 | 541 | 921 | 1,487 | 35.7 | 13.6 | 381.5 | 258.5 | 55.6% | 62.8% |
| 50% | 4,889 | 589 | 983 | 1,595 | 33.3 | 12.5 | 362.2 | 272.1 | 58.5% | 65.8% |
| 60% | 5,268 | 628 | 1,027 | 1,676 | 31.6 | 11.7 | 350.1 | 282.4 | 60.6% | 68.1% |
| 70% | 5,578 | 658 | 1,058 | 1,736 | 30.5 | 11.2 | 341.1 | 288.1 | 62.1% | 69.4% |
| 80% | 5,840 | 684 | 1,087 | 1,790 | 29.4 | 10.8 | 330.2 | 293.9 | 63.4% | 70.6% |
| 90% | 6,082 | 708 | 1,108 | 1,834 | 28.5 | 10.3 | 325.2 | 299.2 | 64.5% | 71.8% |

| | | | | | | | | | | |
|---|---|---|---|---|---|---|---|---|---|---|
| 100% | 6,290 | 732 | 1,123 | 1,872 | 27.8 | 10.0 | 319.6 | 303.8 | 65.3% | 72.8% |
| FIT 50-75, 1 | | | | | | | | | | |
| 10% | 2,182 | 138 | 302 | 497 | 64.3 | 27.6 | 572.3 | 101.9 | 19.9% | 24.6% |
| 20% | 4,183 | 243 | 510 | 797 | 53.9 | 22.0 | 518.4 | 166.1 | 32.9% | 40.0% |
| 30% | 6,049 | 331 | 666 | 1,032 | 46.6 | 18.2 | 475.9 | 210.7 | 42.0% | 50.4% |
| 40% | 7,792 | 408 | 787 | 1,223 | 41.2 | 15.5 | 440.6 | 241.0 | 48.7% | 57.6% |
| 50% | 9,432 | 475 | 883 | 1,383 | 37.1 | 13.6 | 408.2 | 262.1 | 53.8% | 62.8% |
| 60% | 10,982 | 537 | 963 | 1,521 | 33.9 | 12.2 | 384.7 | 280.0 | 57.7% | 66.7% |
| 70% | 12,456 | 593 | 1,029 | 1,641 | 31.4 | 11.1 | 362.8 | 291.6 | 60.9% | 69.6% |
| 80% | 13,873 | 645 | 1,085 | 1,748 | 29.2 | 10.2 | 343.7 | 302.2 | 63.6% | 72.1% |
| 90% | 15,226 | 692 | 1,133 | 1,842 | 27.5 | 9.6 | 326.3 | 309.1 | 65.7% | 73.8% |
| 100% | 16,510 | 737 | 1,175 | 1,927 | 26.2 | 9.1 | 314.0 | 315.0 | 67.4% | 75.2% |
| mt-sDNA 45-75, 3 | | | | | | | | | | |
| 10% | 2,172 | 274 | 537 | 858 | 53.7 | 22.7 | 487.6 | 159.3 | 33.2% | 38.0% |
| 20% | 3,588 | 430 | 791 | 1,255 | 42.3 | 17.0 | 411.8 | 226.8 | 47.3% | 53.6% |
| 30% | 4,585 | 532 | 934 | 1,492 | 36.4 | 14.1 | 367.9 | 261.8 | 54.7% | 61.4% |
| 40% | 5,332 | 603 | 1,026 | 1,653 | 32.8 | 12.4 | 339.2 | 282.6 | 59.2% | 66.0% |
| 50% | 5,913 | 656 | 1,089 | 1,766 | 30.4 | 11.3 | 319.7 | 295.7 | 62.2% | 69.1% |
| 60% | 6,370 | 699 | 1,133 | 1,851 | 28.8 | 10.6 | 305.3 | 304.9 | 64.1% | 71.0% |
| 70% | 6,748 | 734 | 1,169 | 1,920 | 27.6 | 10.1 | 294.7 | 312.2 | 65.6% | 72.5% |
| 80% | 7,063 | 759 | 1,195 | 1,971 | 26.7 | 9.7 | 287.2 | 317.3 | 66.8% | 73.6% |
| 90% | 7,314 | 782 | 1,218 | 2,017 | 26.1 | 9.4 | 280.2 | 320.8 | 67.5% | 74.4% |
| 100% | 7,576 | 814 | 1,240 | 2,070 | 25.2 | 9.0 | 273.8 | 324.7 | 68.7% | 75.5% |
| FIT 45-75, 1 | | | | | | | | | | |
| 10% | 2,640 | 152 | 348 | 556 | 62.5 | 26.8 | 553.3 | 114.3 | 22.1% | 26.7% |
| 20% | 5,073 | 271 | 583 | 895 | 51.3 | 20.9 | 488.6 | 185.3 | 36.1% | 43.1% |
| 30% | 7,339 | 371 | 753 | 1,155 | 43.7 | 16.9 | 440.0 | 232.3 | 45.6% | 53.8% |
| 40% | 9,470 | 455 | 882 | 1,364 | 38.3 | 14.3 | 400.0 | 263.4 | 52.3% | 60.9% |
| 50% | 11,484 | 530 | 988 | 1,540 | 34.1 | 12.4 | 365.2 | 286.2 | 57.6% | 66.1% |
| 60% | 13,388 | 600 | 1,070 | 1,689 | 30.9 | 11.0 | 338.3 | 303.8 | 61.5% | 70.0% |
| 70% | 15,193 | 664 | 1,140 | 1,822 | 28.5 | 10.0 | 316.0 | 315.5 | 64.5% | 72.7% |
| 80% | 16,953 | 718 | 1,199 | 1,933 | 26.5 | 9.2 | 296.6 | 325.6 | 67.1% | 75.0% |
| 90% | 18,599 | 775 | 1,249 | 2,039 | 24.7 | 8.5 | 278.3 | 333.3 | 69.3% | 76.8% |
| 100% | 20,175 | 827 | 1,293 | 2,135 | 23.4 | 8.0 | 264.4 | 338.6 | 70.9% | 78.1% |

COL, colonoscopy; CRC, colorectal cancer; FIT, fecal immunochemical test; LY, life-years; LYG, life-years gained; mt-sDNA, multitarget stool DNA test.

**Table S11. Deep-C (clinicaltrials.gov identifier, NCT01397747) sensitivity analysis of predicted outcomes per 1000 individuals screened from ages 50–75 compared with no screening.** Individuals were randomly assigned numbers of mt-sDNA (max=9 triennial tests during the screening window) or FIT (max=26 annual tests during the screening window).

| Screening strategy | Randomly Assigned Number of Tests | Total Stool Tests | Total COLs | CRC Cases | CRC Deaths | LY with CRC | LYG | Incremental COL/ Incremental LYG vs FIT | Incidence Reduction | Mortality Reduction |
|---|---|---|---|---|---|---|---|---|---|---|
| mt-sDNA, 50-75 | Up to 1 | 819 | 440 | 66.6 | 28.8 | 597.1 | 73.9 | 6.2 | 17.1% | 21.3% |
| FIT, 50-75 | Up to 1 | 813 | 257 | 73.0 | 32.1 | 628.2 | 44.4 | | 9.1% | 12.3% |
| mt-sDNA, 50-75 | Up to 5 | 3,527 | 1,303 | 39.8 | 15.3 | 438.7 | 223.7 | 7.2 | 50.4% | 58.2% |
| FIT, 50-75 | Up to 5 | 3,702 | 743 | 55.1 | 22.1 | 557.1 | 145.6 | | 31.5% | 39.8% |
| mt-sDNA, 50-75 | Up to 9 | 6,057 | 1,832 | 28.5 | 10.2 | 327.1 | 298.5 | 7.9 | 64.5% | 72.0% |
| FIT, 50-75 | Up to 9 | 6,255 | 1,072 | 45.0 | 17.1 | 497.4 | 201.8 | | 44.0% | 53.4% |

COL, colonoscopy; CRC, colorectal cancer; FIT, fecal immunochemical test; LY, life-years; LYG, life-years gained; mt-sDNA, multitarget stool DNA test.

**Figure S1. Age-specific risks of complications from colonoscopy with polypectomy used in the analysis.** Complications include serious gastrointestinal events, other gastrointestinal events, and cardiovascular events. We assume that the only harms from screening arise from a colonoscopy with polypectomy, whether it be for screening, follow-up, or surveillance, or for the diagnosis of a symptomatic cancer. We assume no risk of harms from stool-based tests, nor from bowel preparation.[5] The risks of complications from colonoscopy are from an analysis by van Hees et al.,[6] which was an extension of an analysis by Warren et al.[7] In those studies, colonoscopy without polypectomy was not associated with an excess risk of complications, relative to a matched control group that did not have colonoscopy. Reproduced and adapted with permission from Knudsen et al, 2016.[3]

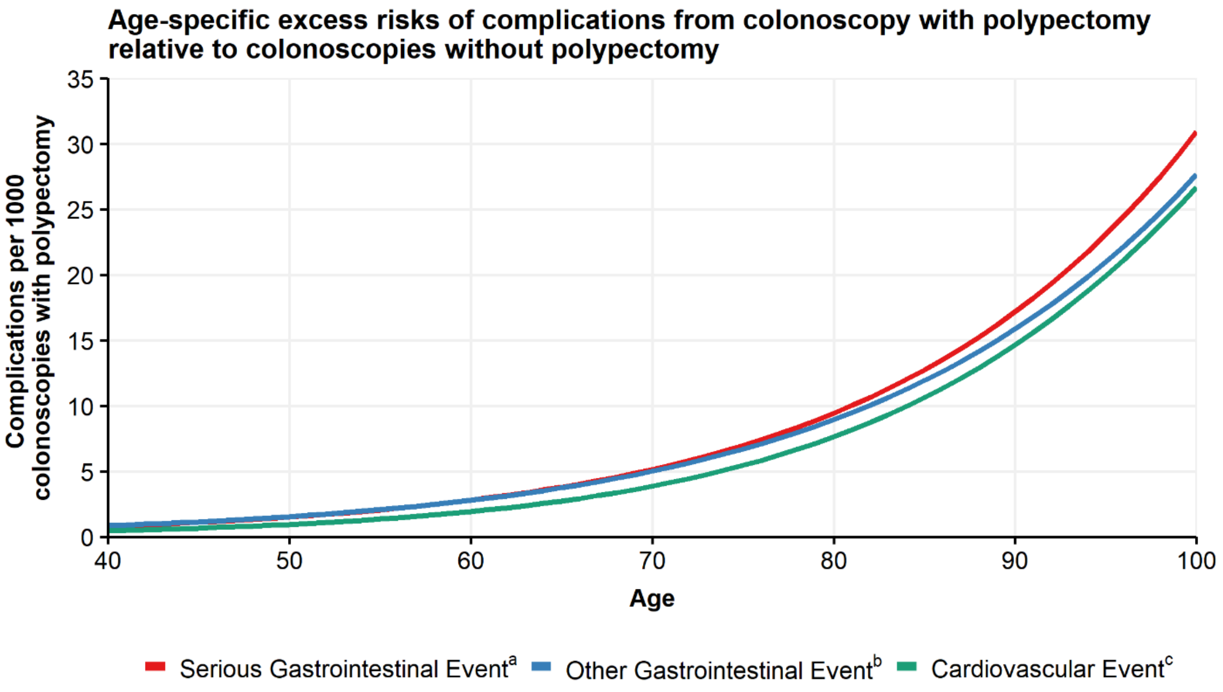


[a] Perforations, gastrointestinal bleeding or transfusions. Excess risk per colonoscopy with polypectomy = $1/[\exp(9.27953 - 0.06105 \times \text{Age}) + 1] - 1/[\exp(10.78719 - 0.06105 \times \text{Age}) + 1]$.

[b] Paralytic ileus, nausea and vomiting, dehydration, abdominal pain. Excess risk per colonoscopy with polypectomy = $1/[\exp(8.81404 - 0.05903 \times \text{Age}) + 1] - 1/[\exp(9.61197 - 0.05903 \times \text{Age}) + 1]$.

[c] Myocardial infarction or angina, arrhythmias, congestive heart failure, cardiac or respiratory arrest, syncope, hypotension, or shock. Excess risk per colonoscopy with polypectomy = $1/[\exp(9.09053 - 0.07056 \times \text{Age}) + 1] - 1/[\exp(9.38297 - 0.07056 \times \text{Age}) + 1]$.

**Figure S2. Percent difference in predicted life-years gained (LYG) per 1000 individuals by adherence rate for triennial multitarget stool DNA (mt-sDNA) versus annual fecal immunochemical test (FIT) in individuals free of diagnosed colorectal cancer at age 40 and screened between ages 45–75 years.** White boxes indicate <10% difference between tests. Light gray boxes indicate ≥10% positive difference with mt-sDNA versus FIT. Dark gray boxes indicate ≥10% negative difference with mt-sDNA versus FIT. Outlined box indicates base-case imperfect adherence rates.

| | Screening Age 45-75 Years | | | | | | | | | | |
|---|---|---|---|---|---|---|---|---|---|---|---|
| **Adherence Rate** | **10** | **20** | **30** | **40** | **50** | **60** | **70** | **80** | **90** | **100** | **Annual FIT LYG/1000 Individuals** |
| **10** | 37.1% | 100.0% | 132.1% | 151.9% | 164.2% | 173.5% | 180.5% | 183.8% | 187.7% | 191.3% | 110.2 |
| **20** | -16.2% | 22.2% | 41.7% | 53.8% | 61.4% | 67.0% | 71.3% | 73.3% | 75.7% | 77.9% | 180.4 |
| **30** | -34.3% | -4.1% | 11.2% | 20.7% | 26.6% | 31.0% | 34.4% | 36.0% | 37.9% | 39.6% | 229.9 |
| **40** | -42.5% | -16.1% | -2.7% | 5.6% | 10.8% | 14.7% | 17.6% | 19.0% | 20.7% | 22.1% | 262.7 |
| **50** | -47.3% | -23.1% | -10.7% | -3.1% | 1.6% | 5.2% | 7.9% | 9.2% | 10.7% | 12.0% | 286.5 |
| **60** | -50.3% | -27.6% | -16.0% | -8.8% | -4.3% | -1.0% | 1.6% | 2.8% | 4.2% | 5.5% | 304.3 |
| **70** | -52.4% | -30.6% | -19.5% | -12.6% | -8.4% | -5.1% | -2.7% | -1.6% | -0.2% | 1.0% | 317.6 |
| **80** | -53.9% | -32.7% | -22.0% | -15.3% | -11.1% | -8.0% | -5.7% | -4.6% | -3.2% | -2.1% | 327.6 |
| **90** | -54.9% | -34.2% | -23.6% | -17.1% | -13.0% | -10.0% | -7.7% | -6.6% | -5.3% | -4.1% | 334.7 |
| **100** | -55.6% | -35.3% | -24.9% | -18.5% | -14.6% | -11.6% | -9.3% | -8.2% | -7.0% | -5.8% | 340.7 |
| **Triennial mt-sDNA LYG/1000 Individuals** | 151.1 | 220.4 | 255.7 | 277.5 | 291.1 | 301.3 | 309.0 | 312.7 | 317.0 | 320.9 | |

**Figure S3. A) Predicted life-years gained (LYG), B) CRC-related incidence and mortality reduction, and C) total stool tests and colonoscopies (COL) per 1000 individuals screened from ages 45–75 compared with no screening assuming perfect (100%) adherence rates to annual FIT, annual HSgFOBT, and triennial mt-sDNA or base-case imperfect adherence (40% FIT vs 34% HSgFOBT vs 70% mt-sDNA).**

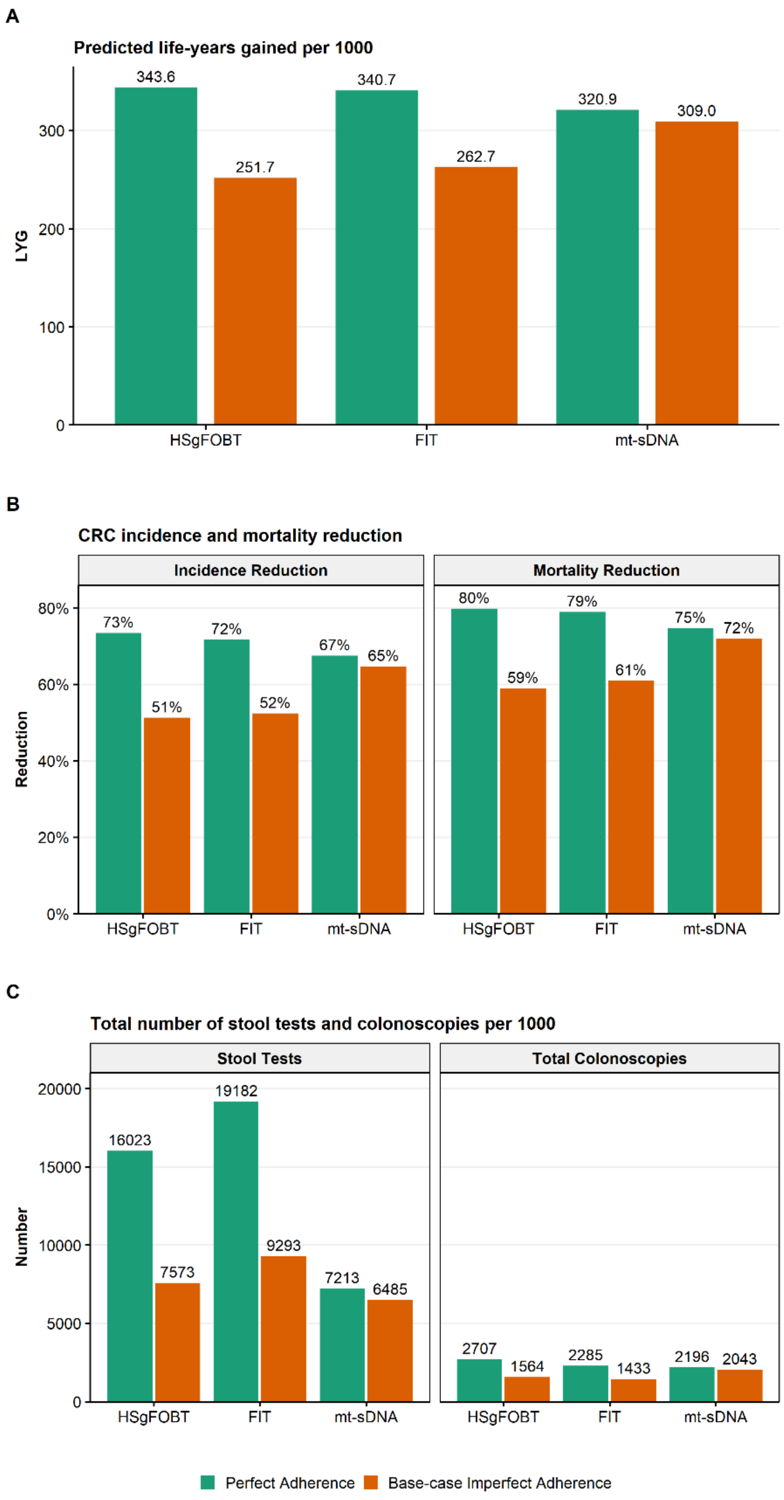

**Figure S4. Predicted life-years gained (LYG) by total stool tests per 1000 individuals screened from ages 45–75 compared with no screening.** Individuals were randomly assigned numbers of multitarget stool DNA (mt-sDNA; max=11 triennial tests during the screening window) or fecal immunochemical tests (FIT; max=31 annual tests during the screening window). The line indicates equivalent LYG with up to 11 mt-sDNA tests.

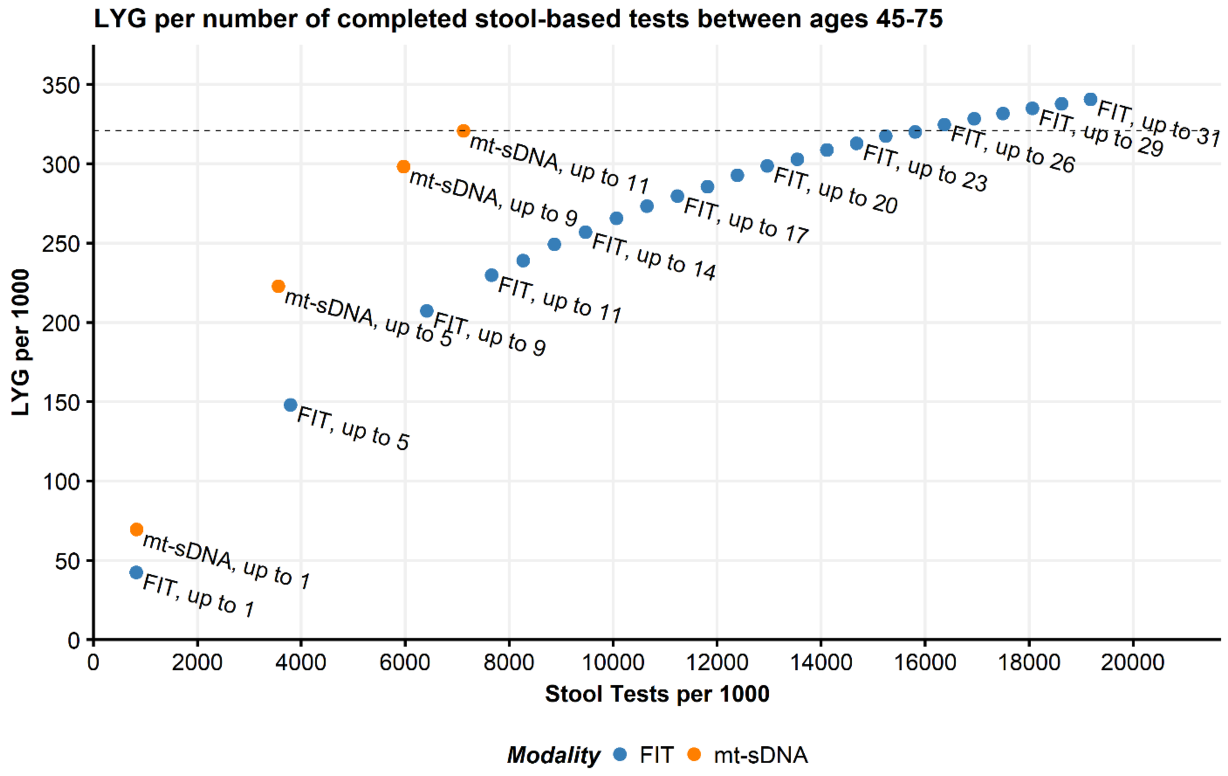

**Figure S5. Deep-C (clinicaltrials.gov identifier, NCT01397747) sensitivity analysis of percent difference in predicted life-years gained (LYG) per 1000 individuals by adherence rate for triennial multitarget stool DNA (mt-sDNA) versus annual fecal immunochemical test (FIT) in individuals free of diagnosed colorectal cancer at age 40 and screened between ages 50–75 years.** White boxes indicate <10% difference between tests. Light gray boxes indicate ≥10% positive difference with mt-sDNA versus FIT. Dark gray boxes indicate ≥10% negative difference with mt-sDNA versus FIT. Outlined box indicates base-case imperfect adherence rates.

**Screening Age 50-75 Years**

| Adherence Rate | 10 | 20 | 30 | 40 | 50 | 60 | 70 | 80 | 90 | 100 | Annual FIT LYG/1000 Individuals |
|---|---|---|---|---|---|---|---|---|---|---|---|
| **10** | 38.3% | 100.3% | 133.6% | 153.8% | 167.2% | 177.3% | 182.9% | 188.6% | 193.8% | 198.3% | 101.9 |
| **20** | -15.2% | 22.8% | 43.3% | 55.7% | 63.8% | 70.0% | 73.5% | 77.0% | 80.2% | 82.9% | 166.1 |
| **30** | -33.1% | -3.2% | 12.9% | 22.7% | 29.1% | 34.0% | 36.7% | 39.5% | 42.0% | 44.2% | 210.7 |
| **40** | -41.5% | -15.3% | -1.3% | 7.3% | 12.9% | 17.2% | 19.6% | 22.0% | 24.2% | 26.1% | 241.0 |
| **50** | -46.2% | -22.2% | -9.2% | -1.4% | 3.8% | 7.8% | 9.9% | 12.1% | 14.2% | 15.9% | 262.1 |
| **60** | -49.7% | -27.1% | -15.0% | -7.7% | -2.8% | 0.9% | 2.9% | 5.0% | 6.9% | 8.5% | 280.0 |
| **70** | -51.7% | -30.0% | -18.4% | -11.3% | -6.7% | -3.2% | -1.2% | 0.8% | 2.6% | 4.2% | 291.6 |
| **80** | -53.4% | -32.5% | -21.3% | -14.5% | -10.0% | -6.6% | -4.7% | -2.7% | -1.0% | 0.5% | 302.2 |
| **90** | -54.4% | -34.0% | -23.0% | -16.4% | -12.0% | -8.6% | -6.8% | -4.9% | -3.2% | -1.7% | 309.1 |
| **100** | -55.3% | -35.2% | -24.5% | -17.9% | -13.6% | -10.4% | -8.6% | -6.7% | -5.0% | -3.6% | 315.0 |
| **Triennial mt-sDNA LYG/1000 Individuals** | 140.9 | 204.0 | 237.9 | 258.5 | 272.1 | 282.4 | 288.1 | 293.9 | 299.2 | 303.8 | |

**Figure S6. Deep-C (clinicaltrials.gov identifier, NCT01397747) sensitivity analysis of predicted life-years gained (LYG) by total stool tests per 1000 individuals screened from ages 50–75 compared with no screening.** Individuals were randomly assigned numbers of multitarget stool DNA (mt-sDNA; max=9 triennial tests during the screening window) or fecal immunochemical tests (FIT; max=26 annual tests during the screening window). The line indicates equivalent LYG with up to 9 mt-sDNA tests.

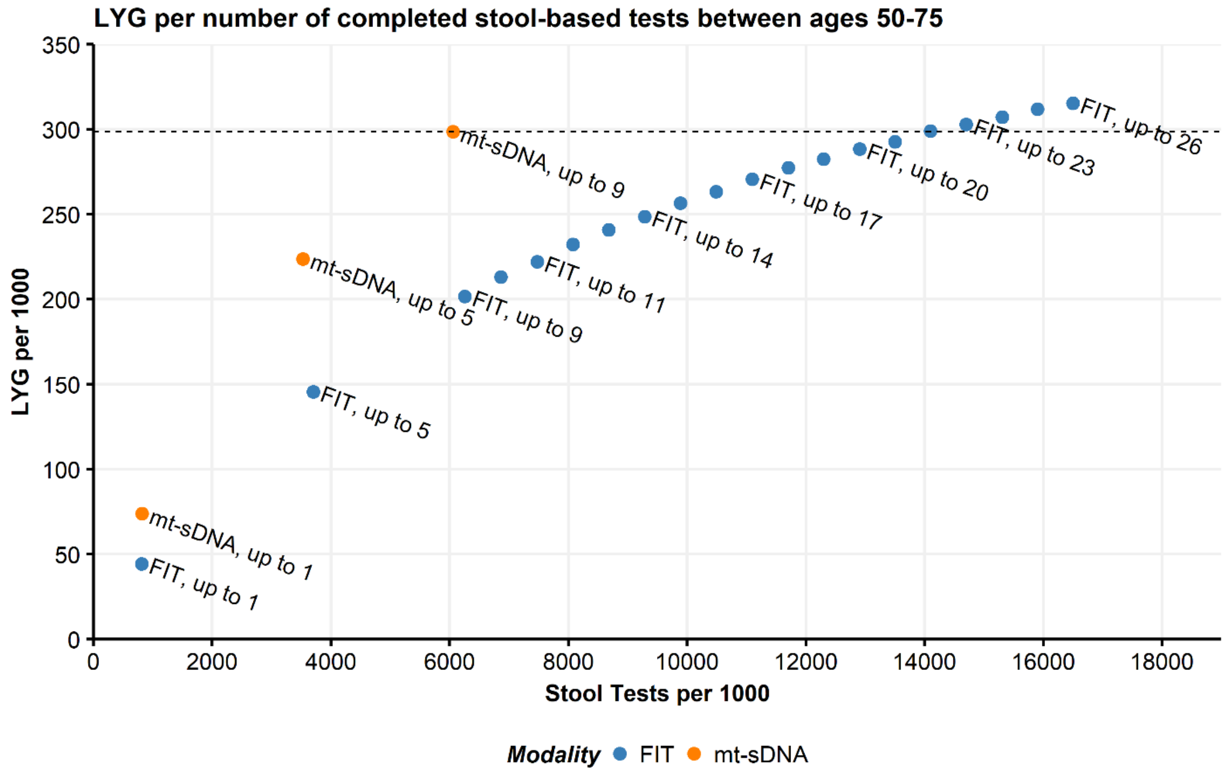

**Figure S7. Sensitivity analysis of life-years gained for individuals 40 years of age with stool-based tests A) number of colonoscopies assuming 50% FIT vs 43% HSgFOBT vs 70% mt-sDNA adherence or B) assuming 60% FIT vs 52% HSgFOBT vs 70% mt-sDNA adherence and by C) patient hours related to the screening process and assuming 50% FIT vs 43% HSgFOBT vs 70% mt-sDNA adherence or D) assuming 60% FIT vs 52% HSgFOBT vs 70% mt-sDNA adherence.** Results shown are per 1000 individuals free of diagnosed colorectal cancer at age 40 and screened starting at age 45 or 50 and ending at age 75 or 80 receiving biennial or triennial mt-sDNA, annual or biennial FIT, and annual or biennial HSgFOBT. NE, near-efficient.

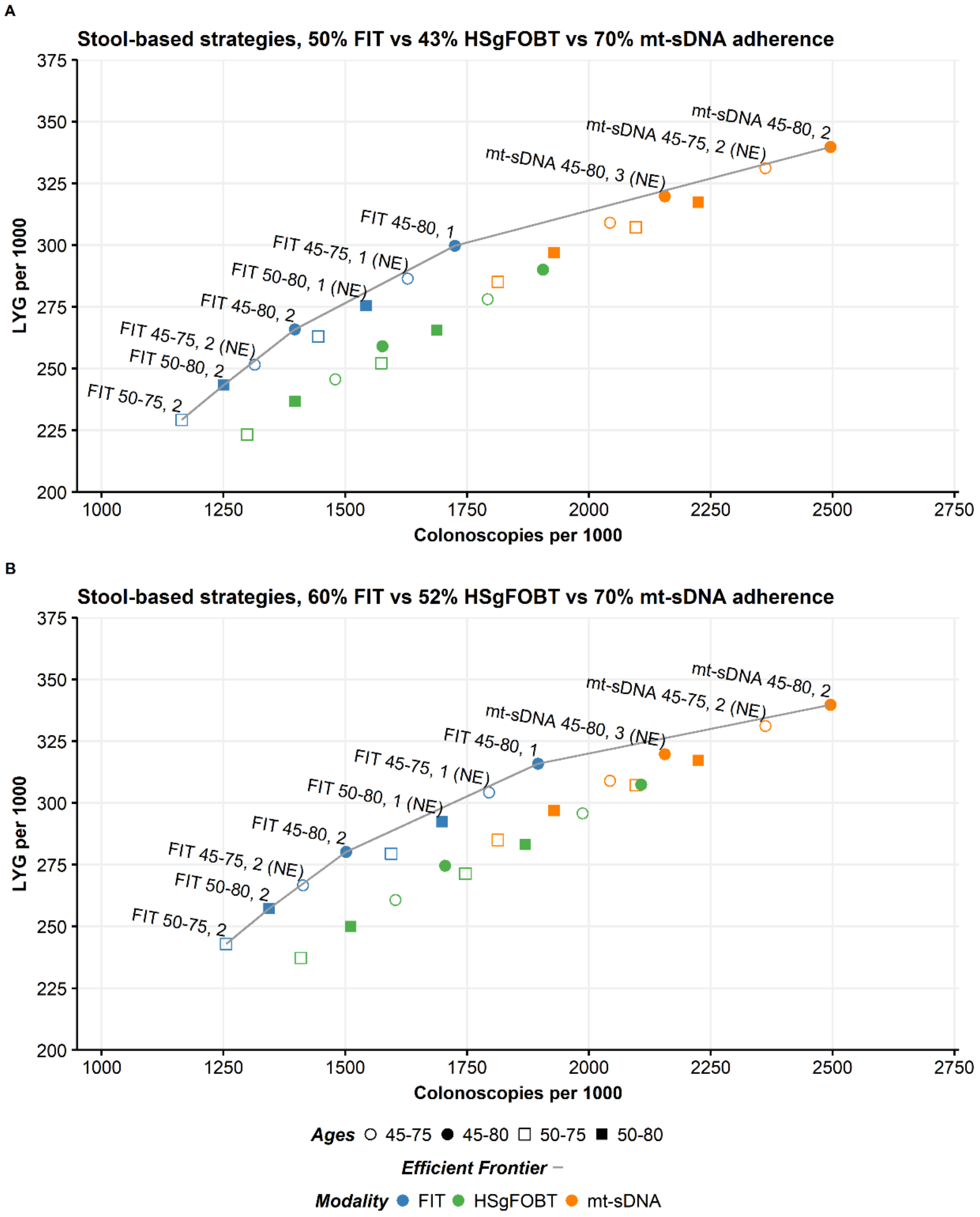
A
Stool-based strategies, 50% FIT vs 43% HSgFOBT vs 70% mt-sDNA adherence
LYG per 1000
Colonoscopies per 1000
375
350
325
300
275
250
225
200
1000
1250
1500
1750
2000
2250
2500
2750
mt-sDNA 45-80, 2
mt-sDNA 45-75, 2 (NE)
mt-sDNA 45-80, 3 (NE)
FIT 45-80, 1
FIT 45-75, 1 (NE)
FIT 50-80, 1 (NE)
FIT 45-80, 2
FIT 45-75, 2 (NE)
FIT 50-80, 2
FIT 50-75, 2
B
Stool-based strategies, 60% FIT vs 52% HSgFOBT vs 70% mt-sDNA adherence
LYG per 1000
Colonoscopies per 1000
375
350
325
300
275
250
225
200
1000
1250
1500
1750
2000
2250
2500
2750
mt-sDNA 45-80, 2
mt-sDNA 45-75, 2 (NE)
mt-sDNA 45-80, 3 (NE)
FIT 45-80, 1
FIT 45-75, 1 (NE)
FIT 50-80, 1 (NE)
FIT 45-80, 2
FIT 45-75, 2 (NE)
FIT 50-80, 2
FIT 50-75, 2
Ages 45-75 45-80 50-75 50-80
Efficient Frontier
Modality FIT HSgFOBT mt-sDNA

C

Stool-based strategies, 50% FIT vs 43% HSgFOBT vs 70% mt-sDNA adherence

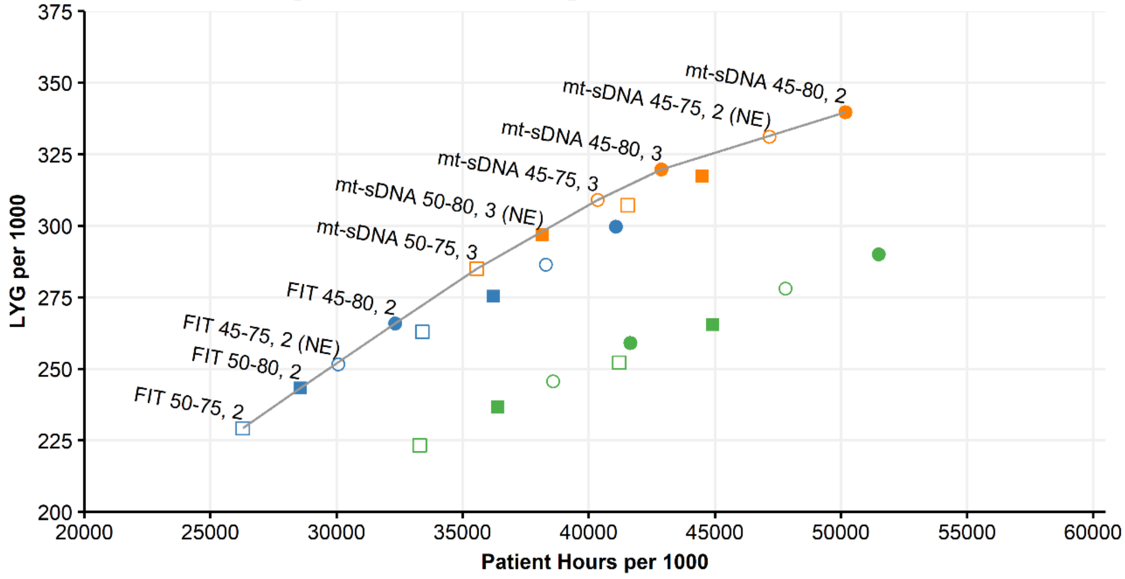


D

Stool-based strategies, 60% FIT vs 52% HSgFOBT vs 70% mt-sDNA adherence

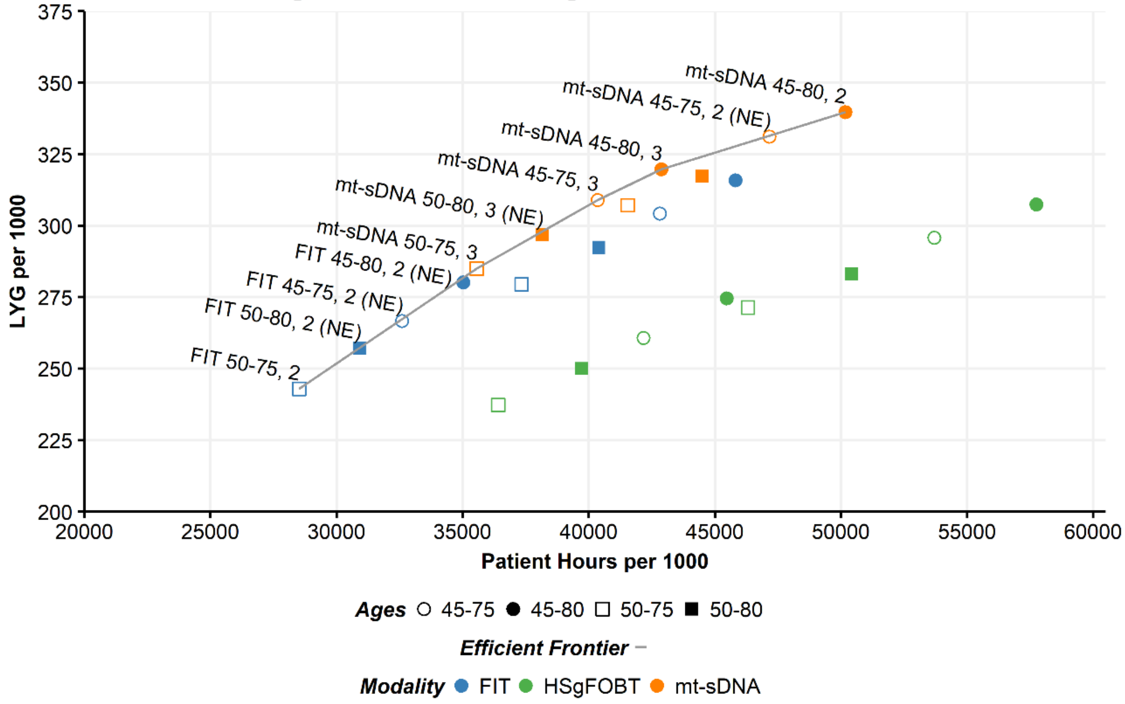

**Figure S8. Life-years gained for individuals 40 years of age with stool-based tests by A) number of colonoscopies or B) patient hours related to the screening process assuming base-case imperfect adherence rates of 40% FIT vs 34% HSgFOBT vs 70% mt-sDNA using the CISNET fixed split adherence approach.** Results shown are per 1000 individuals free of diagnosed colorectal cancer at age 40 and screened starting at age 45 or 50 and ending at age 75 or 80 receiving biennial or triennial mt-sDNA, annual or biennial FIT, and annual or biennial HSgFOBT. NE, near efficient.

A

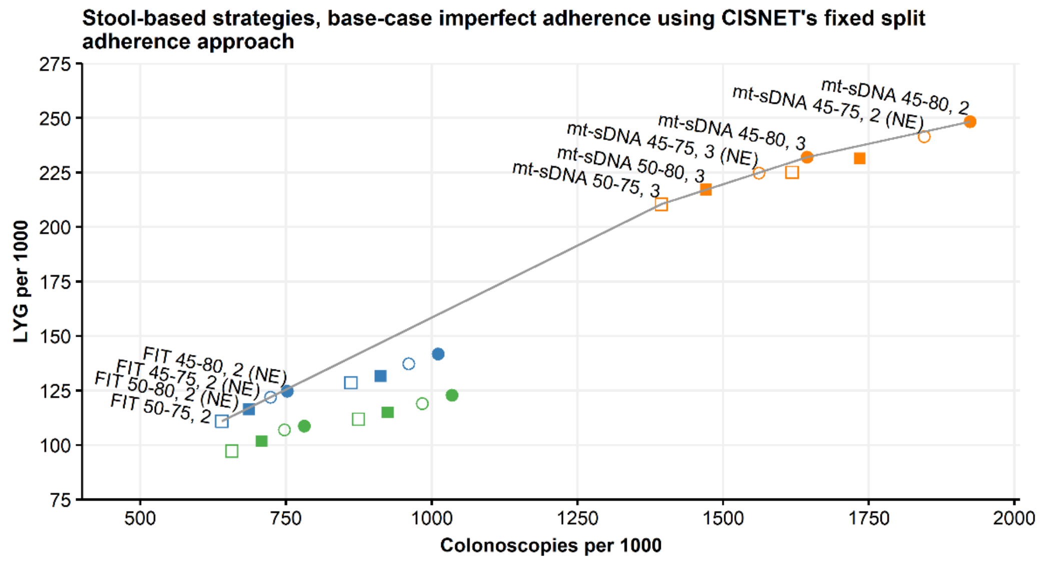


B

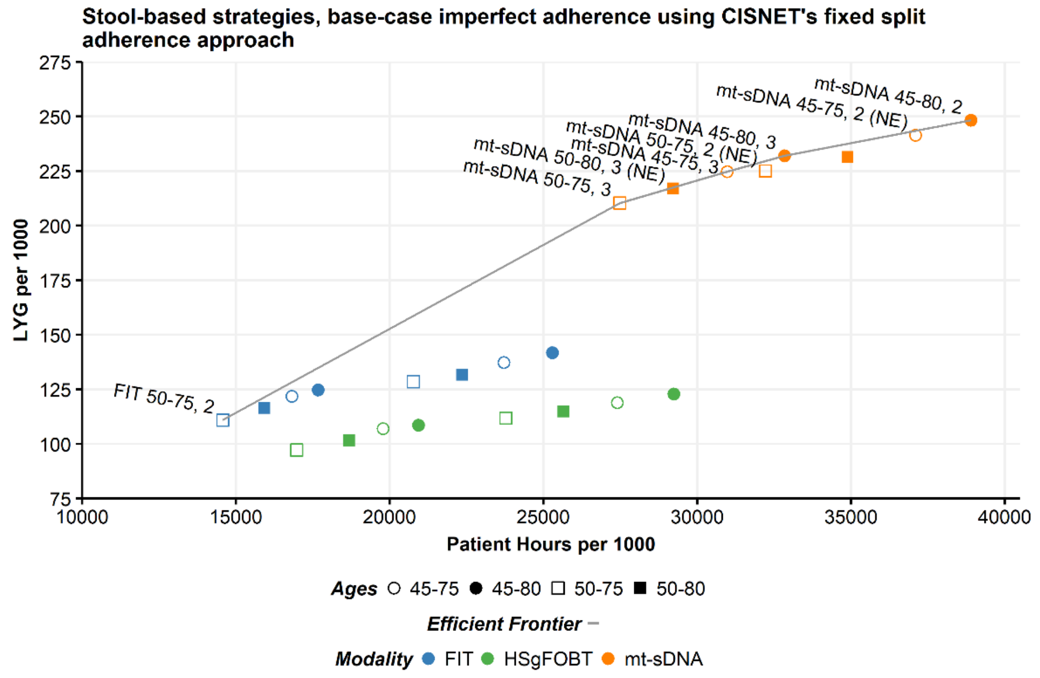
